# UniPTM: Multiple PTM site prediction on full-length protein sequence

**DOI:** 10.1101/2024.08.03.606471

**Authors:** Lingkuan Meng, Jiecong Lin, Ke Cheng, Kui Xu, Hongyan Sun, Ka-Chun Wong

## Abstract

Post-translational modifications (PTMs) enrich the functional diversity of proteins by attaching chemical groups to the side chains of amino acids. In recent years, a myriad of AI models have been proposed to predict many specific types of PTMs. However, those models typically adopt the sliding window approach to extract short and equal-length protein fragments from full-length proteins for model training. Unfortunately, such a subtle step results in the loss of long-range information from distal amino acids, which may impact the PTM formation process. In this study, we introduce UniPTM, a window-free model designed to train and test on natural and full-length protein sequences, enabling the prediction of multiple types of PTMs in a holistic manner. Moreover, we established PTMseq, the first comprehensive dataset of full-length protein sequences with annotated PTMs, to train and validate our model. UniPTM has undergone extensive validations and significantly outperforms existing models, elucidating the influence of protein sequence completeness on PTM. Consequently, UniPTM offers interpretable and biologically meaningful predictions, enhancing our understanding of protein functionally and regulation. The source code and PTMseq dataset for UniPTM are available at https://www.github.com/TransPTM/UniPTM.

## Introduction

Post-translational modifications (PTMs) enrich the functional diversity of proteins by attaching chemical groups or proteins to amino acid side chains. As of now, researchers have identified more than 650 distinct types of PTMs across various proteins, as detailed in UniProt (http://www.uniprot.org/docs/ptmlist.txt).^1^ PTMs encompass phosphorylation, acetylation, methylation, ubiquitination, and others.^2^ The complex processes and wide-ranging functional impacts of PTMs have sparked research interests due to their broad implications in different areas such as enzyme activity modulation,^3^ gene regulation,^4^ and disease treatment.^5^ For example, protein phosphorylation is a reversible PTM where a protein kinase adds a covalently bound phosphate group to an amino acid residue. Phosphorylation changes a protein’s structural conformation, leading to its activation, deactivation, or other modifications in function. ^6^ Around 13,000 human proteins contain sites that can be phosphorylated.^7^ In addition, the N-terminal acetylation of histone lysines plays a major role in gene expression. It opens up the chromatin structure, allowing transcription factors to access the DNA and initiate the transcription process, thereby promoting gene expression.^8^ Moreover, methylation is also one of the most common PTMs, and it regulates functional diversity in the cell. Arginine can undergo methylation in two forms: be methylated once, resulting in monomethylarginine, or dimethylated. In dimethylation, arginine methyltransferases (PRMTs) transfer two methyl groups either to the same nitrogen atom, forming symmetric dimethylarginine, or to different nitrogen atoms at the end of the arginine residue, resulting in asymmetric dimethylarginine.^9^

Because of the countless possible combinations of amino acid residues, enzyme activities, and target sites, there is a sheer number of PTMs, each adding to the variety of protein sequences. Consequently, it is challenging to rapidly and affordably identify the modifications essential for the specific functions of each protein. Thanks to the flourishing of high-throughput mass spectrometry (MS), the field of proteomics has been significantly enriched, resulting in the identification of numerous PTM sites and the emergence of various databases. These data have fostered the development of numerous computational tools for PTM site prediction. Here is a brief literature review on related works: MusiteDeep, the pioneering deep learning-based tool for PTM prediction, offers general predictions for phosphosites as well as kinase-specific phosphosite predictions for five kinase families, each with over 100 known substrates. In 2020, they enhanced the tool by launching a web server that allows for the prediction and visualization of multiple types of protein PTM sites. This server can predict multiple types of PTM sites simultaneously at the amino acid level.^10^ Also, PTM-CMGMS incorporates not only traditional protein sequence data but also structural information predicted by AlphaFold^11^ to predict crotonylation, succinylation, and nitrosylation PTM sites.^12^ In addition, given the sequential nature of protein sequences, protein language models (pLMs) have been developed to learn insightful representations of proteins using the vast amount of unlabeled sequences available in protein databases. Peng et al. trained PTM-Mamba, a pLM that utilizes a bidirectional gated Mamba block to represent both wild-type and post-translationally modified protein sequences, effectively capturing PTM-specific effects within its latent space.^13^ Moreover, Gutierrez et al. proposed a deep learning model, Sitetack, for predicting multiple types of PTM sites. ^14^ This model encodes known PTMs as separate amino acids in the protein sequence during the data representation stage and then inputs the embedded data into a convolutional neural network (CNN) architecture. Sitetack outperforms MusiteDeep in predicting 11 types of PTM sites, particularly in terms of the area under the precision-recall curve (AUPRC).

However, those models typically use the sliding window approach ^15^ to extract short equal-length protein fragments from the full-length protein for model training. Indeed, the sliding window data processing method cannot capture the influence of distal amino acids on the PTM site. Since real proteins are three-dimensionally folded, distal amino acids can be folded and closely positioned next to each PTM site of interest. These models are only able to consider sequence elements up to 35 amino acids away from the target amino acids^16^ because the sliding window strategy significantly limits sequence data length; for example, phosphorylase kinase has complex binding requirements because all residues spatially near the phosphorylatable site are crucial for phosphorylation. Additionally, higher-order structures are important for the specificity and efficiency of the phosphorylation process.^17^ Thus, capturing the information flow between distal residues is a promising path for further studies. To address this, we developed UniPTM (Figure 1), a window-free transformer-based model designed to predict nine types of PTM site, taking into account real full-length proteins with the capability to annotate multiple sites even for the same PTM type. Each amino acid in the full-length protein is individually embedded into 1280-dimensional vectors using the pre-trained ESM-2 pLM.^18^ This per-residue embedding ensures that each amino acid is taken into consideration. Before these embeddings are processed by encoding layer, they are reduced to 256 dimensions with a convolutional layer. Subsequently, they are fed into an eight-head transformer encoder for feature extraction, followed by a binary classification module in three fully-connected layers. To the best of our knowledge, there is not any existing dataset dedicated to directly offer labeled PTM sites on full-length proteins. To address this gap, we developed PTMseq, a dataset comprised of 12,203 full-length protein sequences annotated with nine distinct types of PTMs. This dataset features a total of 34,514 PTM sites, providing a compiled resource for training and testing the UniPTM model.

**Figure 1:**
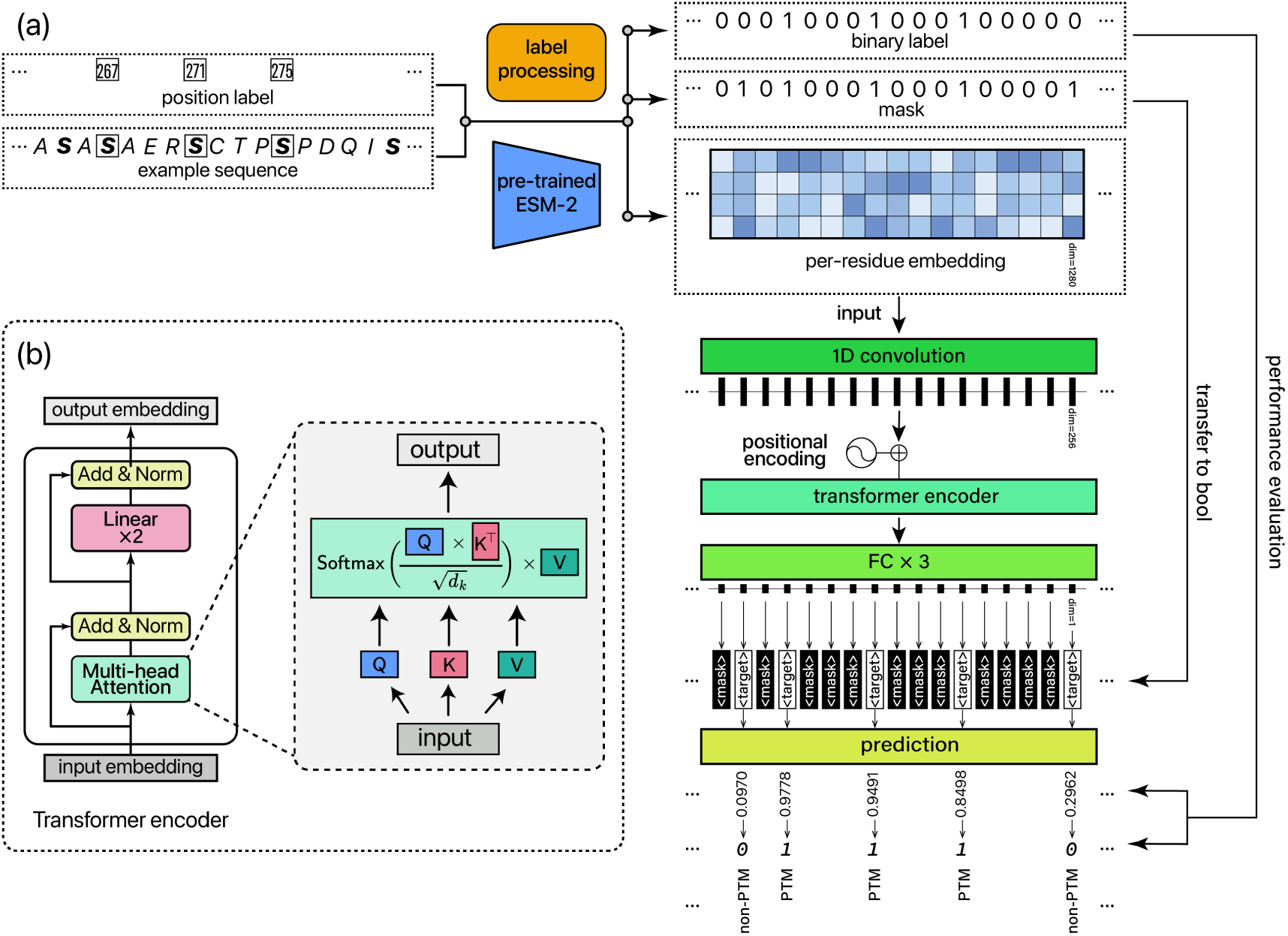
(a) The overall architecture of UniPTM. Phosphorylated full-length protein sequence (antiviral protein UniProt ID: Q7Z2W4 is used as example) is input to the model, the sequence and position label are processed separately by pre-trained ESM-2 protein language model and label generator to yield a 1280-dimensional embedding along with binary label and mask. The embedding is first inputted into a convolutional layer to reduce its dimensionality to 256 dimensions, followed by positional encoding before being fed into a transformer encoder. The transformer-encoded features are then passed through three fully connected (FC) layers, resulting in a one-dimensional array of predictions that match the length of the input sequence. After filtering through the mask, predictions for the target sites are classified (with a threshold of 0.5) and undergo performance evaluation by comparison with the true binary label. A value of 1 at position indicates a PTM site, whereas a value of 0 indicates non-PTM at that site. (b) The components of a transformer encoder layer. Features are processed through the multi-head scaled dot-product attention layer followed by linear layers. Residual connections and layer normalization are implemented after those two steps.

## Methods

### Data curation

Given the lack of existing databases or literature providing datasets annotated with PTM sites on full-length proteins, we were compelled to develop the PTMseq dataset from scratch to align with our study. The creation of the PTMseq dataset, as depicted in Figure 2a, involved three main stages: data collection, data cleaning and categorization, and dataset construction.

**Figure 2:**
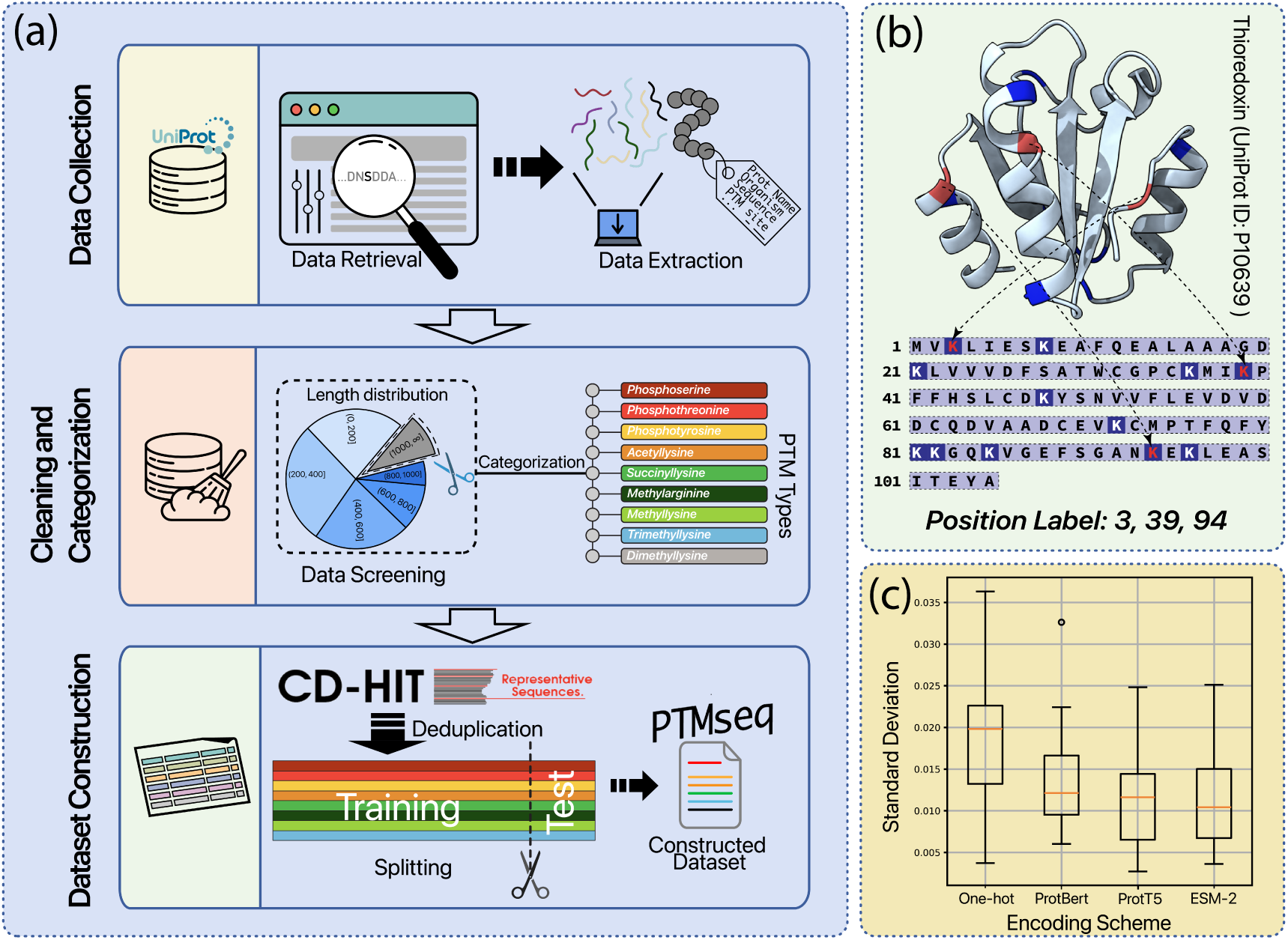
PTMseq dataset construction. (a) Workflow for the creation of the dataset. (b) Example data of acetylated full-length sequence and its position label. (c) Standard deviation of accuracy for different encoding methods across nine PTM subsets.

Initially, we harvested experimentally verified full-length protein sequences from the UniProt database.^1^ By applying filters for *reviewed* status and *proteins with modified residues*, we identified 45,934 full-length protein sequences from over 200 million entries in UniPro-tKB. We selected and downloaded data customized to our specific needs, including columns for entry name, gene name, protein length, sequence, and modified residues. In the subsequent step, we sorted the downloaded full-length proteins by sequence length and removed those exceeding 1000 amino acids, which constituted only 8.76% of all sequences (Figure S1). We then processed the remaining 22,428 sequences by extracting *modified residue* information, which identifies the PTM sites on each sequence. Sequences may appear multiple times, because each sequence can be modified in different ways depending on the specific PTM involved. From this, we organized all the full-length sequences into 41 distinct PTM subsets. Many of these categories were either too small or highly redundant, leading us to select sequences annotated with nine specific PTM types to establish the dataset: Phospho-serine, Phosphothreonine, Phosphotyrosine, N6-acetyllysine, N6-succinyllysine, Omega-N-methylarginine, N6-methyllysine, N6,N6,N6-trimethyllysine, and N6,N6-dimethyllysine. For clarity throughout this paper, these nine PTMs are abbreviated as Phosphoserine, Phosphothreonine, Phosphotyrosine, acetyllysine, succinyllysine, methylarginine, methyllysine, trimethyllysine, and dimethyllysine. In the final step, we employed the CD-HIT tool^19^ to remove redundant sequences from each subset of the nine PTMs, resulting in 12,203 unique full-length protein sequences. The deduplication details of each PTM subset are shown in Figure S2. These sequences were then randomly split into training and independent test sets in a 4:1 ratio, with the division stratified by sequence length to ensure uniform length distribution across both sets. To validate the robustness of our dataset division, we conducted a 5-fold cross-validation (CV) on the entire dataset using four encoding schemes: one-hot, ProtBert, ProtT5, and ESM-2 (Table S1). Each of the nine PTM subsets was encoded using these four methods and evaluated with our UniPTM model through 5-fold CV. The standard deviations of the CV results across all nine subsets and encoding methods were less than 0.02, which are less than 2% of the mean (Figure 2c). Therefore, we consider this dataset division method to be equitable and effective. This methodical approach led to a dataset comprising of 12,203 full-length protein sequences with 34,514 distinct PTM sites labeled.

Each data label, an integer or a single-digit array, represents one or more modified position on the protein sequence, which we refer to as the ”position label” (as exemplified by the thioredoxin protein in Figure 2b). The PTMseq dataset’s statistics are detailed in Table 1, and the dataset is maintained on our GitHub page, with regular updates every three months to ensure it continues to serve as a valuable resource for the community in this field.

**Table 1:** Statistics for PTMseq dataset.

| PTM type | Training |  |  | Independent Testing |  |  |
| --- | --- | --- | --- | --- | --- | --- |
|  | Positive Site | Negative site | Sequence Count | Positive Site | Negative site | Sequence Count |
| Phosphoserine | 17,105 | 178,061 | 4,810 | 4,135 | 44,381 | 1,203 |
| Phosphothreonine | 4,078 | 56,055 | 2,216 | 1,018 | 13,409 | 554 |
| Phosphotyrosine | 2,091 | 14,822 | 1,047 | 513 | 3,502 | 262 |
| Acetyllysine | 2,661 | 28,166 | 998 | 720 | 7,095 | 250 |
| Succinyllysine | 775 | 5,087 | 208 | 212 | 1,303 | 52 |
| Methylarginine | 511 | 8,523 | 255 | 135 | 2,196 | 64 |
| Methyllysine | 200 | 2,901 | 98 | 50 | 914 | 25 |
| Trimethyllysine | 125 | 1,645 | 76 | 34 | 296 | 20 |
| Dimethyllysine | 123 | 1,349 | 52 | 28 | 333 | 13 |
| **Total** | **27,669** | **296,609** | **9,760** | **6,845** | **73,429** | **2,443** |

### Embedding full-length sequences using pre-trained pLMs

Protein language models (pLMs) predict protein behaviors by treating amino acids as words in a sentence, applying natural language processing (NLP) techniques from large language models. This approach has rapidly gained popularity in recent years. ProtBert,^20^ ProtT5,^21^ and ESM-2^18^ have been pre-trained on more than 200 million natural protein sequences, producing latent embeddings that effectively capture relevant physicochemical and functional information. In this study, we utilize these three state-of-the-art pLMs as fixed feature extractors to extract per-residue embeddings from protein sequences, facilitating PTM site prediction on full-length sequences. In addition to these three methods, one-hot encoding has also been implemented as a control to benchmark the capabilities of pre-trained models. ProtBert is an auto-encoder model developed under the self-supervised BERT architecture, and is trained on the UniRef100 corpus, which contains 217 million protein se-quences.^22,23^ ProtBert employs bidirectional training of transformers, which is integrated into both the encoder and decoder architecture. This approach helps in deepening the structural relationships among amino acids. The ProtBert classification model serves as the foundational architecture for predicting protein secondary structures and subcellular locations, utilizing diverse datasets including CASP12, CB513, and DeepLoc. In the work, it was employed as a sequence embedding tool. When a full-length protein sequence is inputted, ProtBert generates a contextually informed embedding with the dimensions L×1024 where L represents the length of the input sequence.

ProtT5 is a top model from the ProtTrans series, utilizing the T5 method. It was trained on the BFD (Big Fantastic Database), which contains 2.5 billion sequences, and fine-tuned on Uniref50, consisting of 45 million sequences.^22^ Positional encoding is tailored for each attention head in the transformer architecture and is consistently shared across all layers in the attention stack. We adopted the pre-trained ProtT5 model (prot t5 xl uniref50) to extract per-residue embeddings from the last hidden layer of the encoder model. This model takes the complete protein sequence as input and outputs an embedding vector of size 1024 for each amino acid on the protein sequence.

ESM-2 is a cutting-edge protein model trained through masked language modeling to predict properties such as stability, folding rate, and binding affinity. It has been trained on 250 million protein sequences along with their associated properties, enabling it to discern the intricate connections between amino acid sequences and the structural properties of proteins. We employ the ’esm2 t33 650M UR50D’ version of ESM-2, which encompasses 6.5 million trainable parameters and is trained on the UniRef50 database, to produce embeddings of size L×1280.

One-hot encoding serves as a baseline method for benchmark control. With this setup, we can explicitly compare the differences in performance between embeddings generated by pre-trained models and traditional methods. Similar to our previous work,^24^ one-hot encoding produces a vector of size 20 for each amino acid residue.

Following the extraction of these embeddings, protein sequences will be processed through 5-fold CV within our UniPTM model to select the best-performing embedding method for further downstream comparative and analytical tasks.

### Binary label and sequence mask generation

For PTM site prediction tasks, we focus on two key information from the PTMseq dataset: the full-length protein sequences and the position labels, where the indices in the position labels indicate PTM sites. We need to generate binary labels and masks based on these two information to obtain positive and negative sample sites. For a given full-length protein sequence, the corresponding binary label and mask are both one-dimensional binary arrays of the same length as the sequence. In the binary label, positions indicated by the position labels are marked as 1, while all other positions are marked as 0. In the mask, positions corresponding to the amino acid under investigation (referred to as the target amino acid) are marked as 1, while all other positions are marked as 0. Each full-length sequence data is varied in sequence length and the number of PTM sites; for instance, the thioredoxin protein data is taken as an example and mentioned in the data curation subsection. Specifically, Figure 3 illustrates its sequence, position label, binary label, and mask. This protein has a sequence of 105 amino acid residues, and its position label is ’3, 39, 94’. The binary label for this sequence is a one-dimensional array of length 105, where positions 3, 39, and 94 are set to 1, indicating they are acetylated sites (positive site samples), and all other positions are set to 0. The mask is also a 105 amino acid-length one-dimensional binary array, where positions corresponding to the lysine (K), the target for acetylation, are set to 1, and all other positions are set to 0. Positions that have a mask value of 1 and a binary label value of 0 are considered negative site samples.

**Figure 3:**
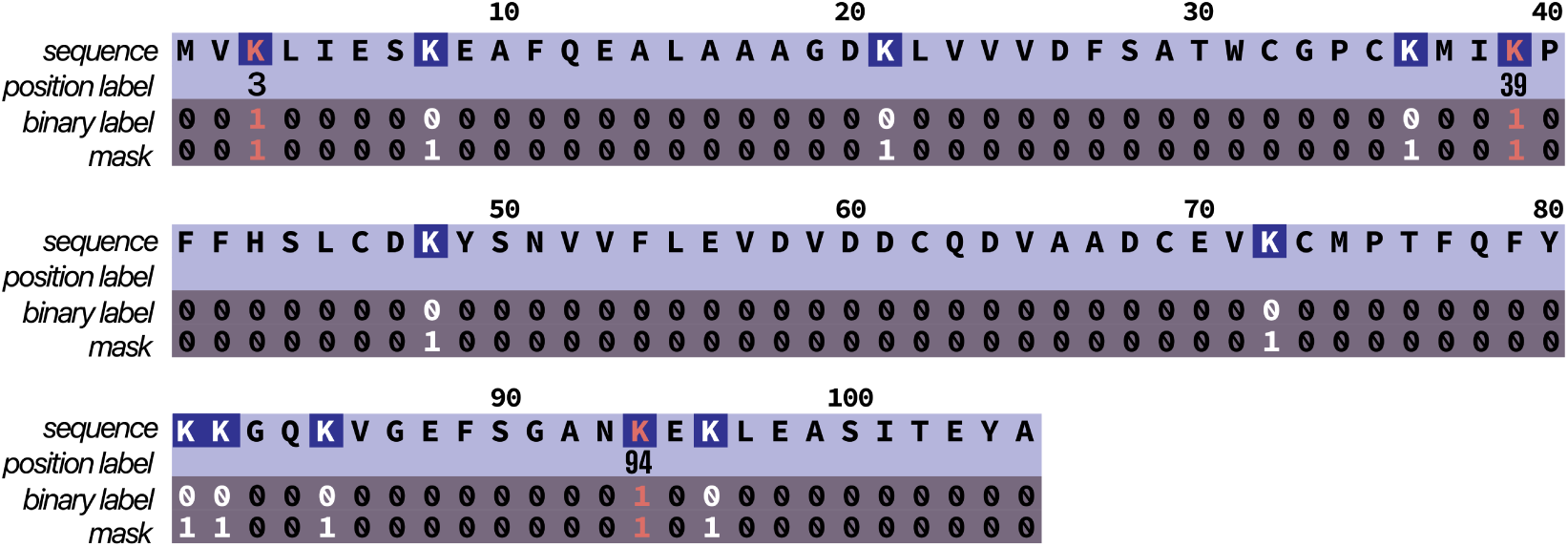
Position labels of the example full-length protein sequence data (thioredoxin protein, UniProt ID: P10639), with corresponding binary label and mask, highlighting acetylated sites and lysine positions.

In summmary, the positive site samples represent lysines that have been acetylated in the sequence, while the negative site samples are lysines that have not been acetylated. As illustrated in Figure 3, there are a total of 3 positive site samples and 9 negative site samples on the example sequence.

### Architecture of UniPTM

The UniPTM architecture is composed of three main components: (1) a shallow convolutional layer, (2) an eight-head transformer encoder, and (3) three fully connected (FC) layers for classification (Figure 1). UniPTM takes as input pLM-embedded full-length protein sequences, as observed in the PTMseq dataset. The shallow convolutional layer initially reduces the spatial dimension of the input embeddings (per-residue) to 256, while the convolutions simultaneously observe amino acids from the current position and adjacent regions. By the convolutional layer, the model can retain essential features while eliminating redundancy, noise, and the risk of overfitting.^25,26^ The transformer encoder has a dimensionality (d model) of 256 and utilizes multi-head attention which captures long-range interactions and contextual information across the entire sequence. pLM-pre-embedded sequences are integrated with positional encoding (as illustrated in Figure 1a) to include information regarding the relative and absolute positions of each feature. ^27^ The calculation of positional encoding is performed by

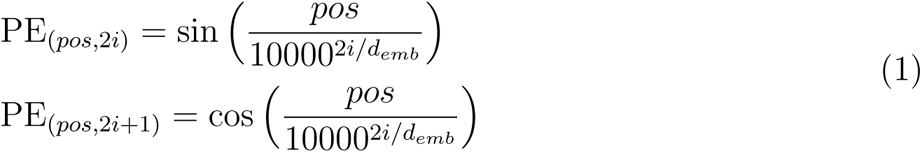

where *pos* represents the feature’s position in the sequence, *i* is the dimension index, and *d*_emb_=256 denotes the embedding dimension. The transformer model is a deep neural network architecture based on the self-attention mechanism (explained in the Supporting Information). Both the attention and MLP components utilize residual connections^28^ and layer normalization.^29^ For each head within the attention layer (see Figure 1b), the input sequence embedding *X* is transformed into the query, key, and value vectors (*Q*,*K*,*V*) by multiplying *X* with the three learnable weight vectors *W_q_*, *W_k_*, and *W_v_*. The scaled dot-product attention, *A*, is computed with the following equation:

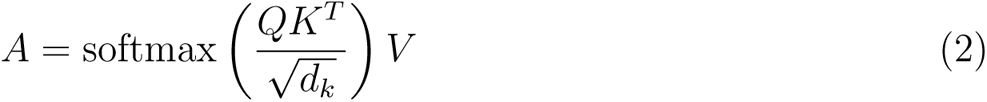

where *d_k_* represents the dimension of *Q* and *K* (as detailed in Supporting Information). In each head, the model initializes the vectors *W_q_*, *W_k_*, and *W_v_* randomly, enabling it to capture the contextual relationships between features across various representational subspaces.^30^ The attentions from all heads are concatenated and subsequently processed through the MLP to produce the projected output embedding, which retains the same dimension as the input embedding. Since the self-attention mechanism integrates information from the entire sequence into each feature embedding, in theory, any single embedding could serve as a representation of the entire sequence. Thus, we adhered to the experimental protocols in related studies^31–33^ by sticking to the output of each target amino acid to calculate the loss during model training.

Finally, the architecture incorporates three FC layers, each utilizing ReLU activation and dropout. These layers sequentially reduce the dimensionality from 256 to a hidden size of 128, ultimately producing a single-dimensional array output for the protein sequence. PTM probability score of each amino acid, denoted as S, is calcualted by:

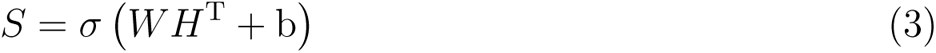

Here, *W ∈* R^1*×*128^ denotes the weight matrix, and *b ∈* R represents the bias term. *H* is the output from the final layer. The sigmoid function is applied to convert this output into a range of (0,1). Hence, the PTM probability scores across the full-length protein sequence are represented as:

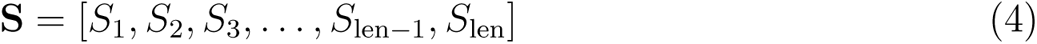

where *len* denotes the length of the sequence, and *S_i_* represents the PTM probability score at the *i*-th position of the sequence. Probability scores greater than 0.5 are classified as PTM sites, while those below 0.5 are classified as non-PTM sites.

### Model training and evaluation

We approached the prediction of PTM sites as a binary classification problem and trained a specialized model for each of the nine PTM types. All these models share the same set of hyperparameters during training. In the training process, the framework accepts full-length protein sequences as input and extracts embeddings from a pre-trained pLM. To enable batch processing of sequences from native proteins, which vary in length, the dataloader was enhanced to pad each sequence embedding, its corresponding binary label, and mask with zeros. This padding ensures that every sequence data within a batch aligns to the length of the longest sequence present in that batch. In addition to the challenge of various sequence lengths, this study faces two other issues: the positions and numbers of positive and negative site samples are not fixed on a sequence; and there is an imbalance between positive and negative site samples. To address these two issues, we customized the binary cross-entropy (BCE) loss function and named it TargetedBCELoss:

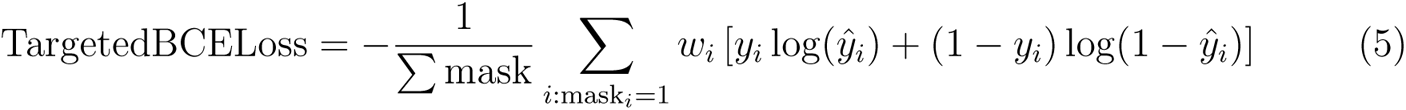

where *y_i_* are the true binary labels, *y̑*_*i*_ are the predicted probabilities; mask *_i_* is a binary mask where the loss is calculated only for indices *i* where mask *_i_*= 1; *w_i_* are the weights applied to each term in the loss calculation, with *w_i_* = pos weight if *y_i_* = 1 and *w_i_* =1 otherwise; ∑ mask denotes the sum of the mask values, which counts the number of elements contributing to the loss, ensuring normalization only over the target amino acid positions.

The UniPTM model was trained to minimize the TargetedBCELoss. To select the best pre-trained pLM, we employed 5-fold stratified cross-validation. In this method, the pre-trained pLM embedded training set is divided into five equal subsets. The model is trained iteratively on four subsets and validated on the fifth. This cycle is rotated and repeated five times, with the average of the validation results from all five iterations to assess performance. Once the best-performing pLM is identified, we use its embedded training set to train the UniPTM model. During training, 10% of the data in the training set is initially set aside as a validation set to monitor training effectiveness at each epoch. If performance ceases to improve after several epochs, early stopping is triggered automatically, and the best model is saved. The trained UniPTM model is then tested on an independent testing subset in the PTMseq dataset. The performance was assessed in five metrics: accuracy, precision, recall, f1 score, and MCC (Matthew’s correlation coefficient), all with a decision probability threshold of 0.5. Additionally, the area under the ROC curve and the area under the precision-recall (PRC) curve served as further performance indicators. This training and testing cycle was repeated five times to compute the average and standard deviation of these metrics. The specific hyperparameters used for the 5-fold CV and independent testing, as well as the mathematical details for the five metrics, are provided in Supporting Information.

## Results

### Performance evaluation of UniPTM

In this section, we evaluate the prediction performance of UniPTM by employing 5-fold cross-validation on the PTMseq dataset’s training set. Figure 4 displays the histograms of AUC, AUPRC, and MCC (Matthews Correlation Coefficient) values, offering a comparative analysis of the performance between ProtBert, ProtT5, and ESM-2 embeddings for full-length sequence data across nine PTM subsets. Accuracy, precision, recall, F1 score, and MCC values, along with AUC and PRC values, are tabulated in Table S3. We first selected AUROC and AUPRC as criteria for selecting the appropriate pLM. AUROC assesses the model’s ability to distinguish between positive and negative site samples, while AUPRC is particularly valuable for reflecting the model’s capability to identify the minority class, which is essential given that our positive site samples are significantly fewer than negative ones. In Figure 4a, the AUROCs for full-length protein sequences embedded via three pLM methods in UniPTM model show similar performance, significantly outperforming the one-hot encoding that used as a negative control. This is expected as one-hot encoding lacks the contextual and structural awareness provided by pLMs. ProtT5 often slightly outperforms ProtBert in terms of AUROC, which might suggest it has better discriminative power for our tasks. ESM-2 shows competitive performance, often close to or slightly better than ProtT5 in some PTMs (e.g., phosphotyrosine in AUROC). In Figure 4b, it remains challenging to distinguish which of the three pLMs performs best in the UniPTM model based on the embedded sequences. This difficulty arises because the highest AUPRC scores vary among the pLMs depending on the type of PTM. For instance, ProtBert achieves the highest AUPRC in the dimethyllysine dataset, while ProtT5 leads in phosphothreonine. Meanwhile, ESM-2 outperforms the other two pLMs in phosphotyrosine and methylarginine. Therefore, we employed another evaluation criterion, MCC, to further attempt to single out the best pLM for UniPTM. MCC is particularly suitable for imbalanced datasets because it provides a more equitable assessment by equally considering all four categories of classification outcomes in the confusion matrix. In Figure 4c, ESM-2 achieves the highest MCC in 7 out of the 9 PTMs examined. As a result, we selected ESM-2 as the embedder for full-length protein sequences to work in conjunction with the UniPTM model for subsequent state-of-the-art comparisons and analyses.

**Figure 4:**
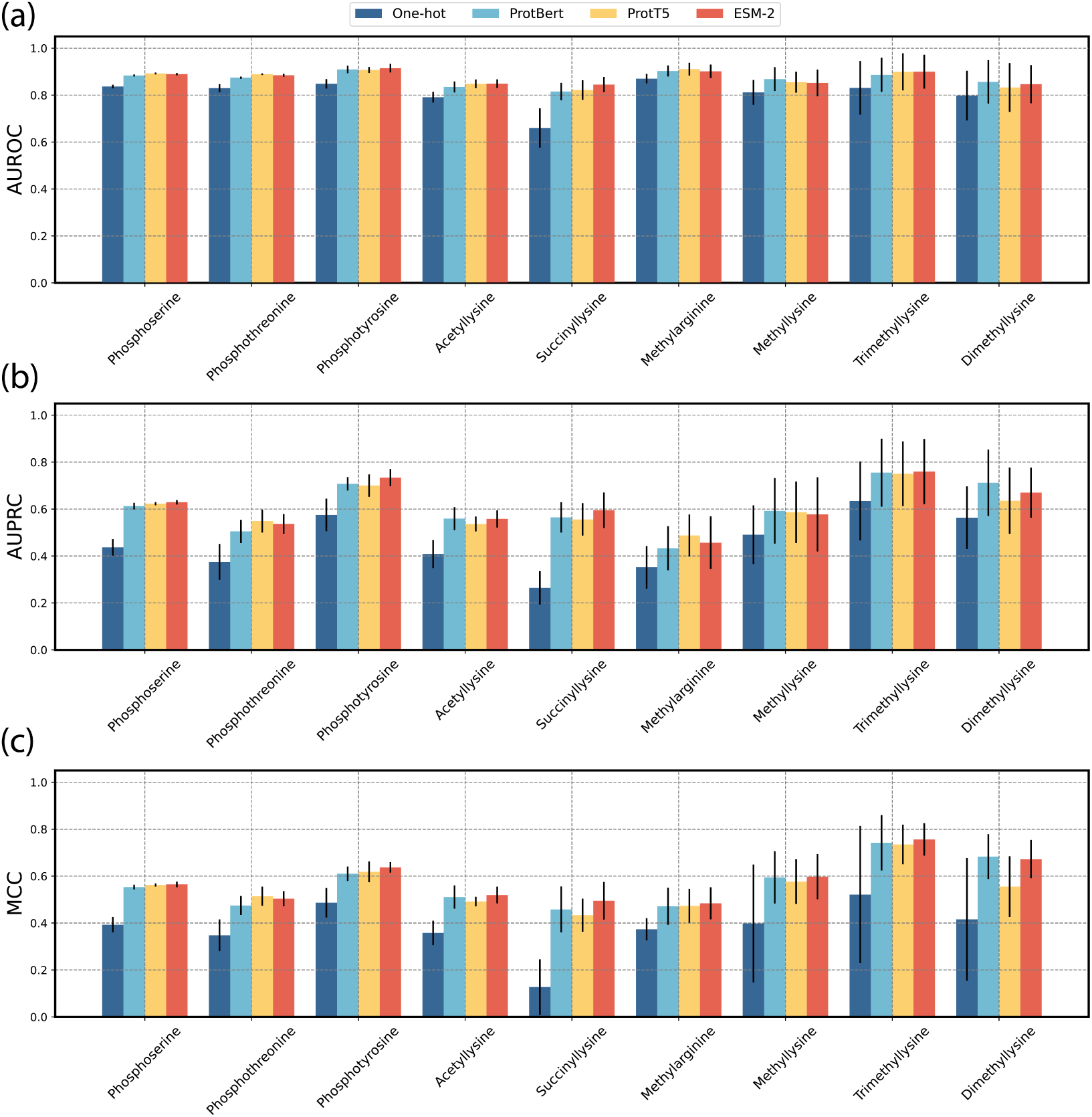
Histogram of 5-fold cross-validation on training sets across 9 types of PTMs. (a) AUROC values for embedding methods: ProtBert, ProtT5, and ESM-2. (b) AUPRC values for the three embedding methods. (b) MCC values for the three embedding methods. one-hot encoding is served as a negative control.

### Comparison with existing methods

To demonstrate the practical performance of PTM site prediction with UniPTM, we compared it to existing tools using the independent testing set of PTMseq. All these comparative tools handle only protein sequence fragments processed using a sliding window strategy, unlike our UniPTM model, which processes unsegmented, variable-length full-length protein sequences. Numerous predictive tools have been developed for each type of PTM. However, due to the unavailability of many of their repositories or web servers, we selected the most widely used and user-friendly tools for comparing each PTM type. Phosphoserine, phospho-threonine, and phosphotyrosine are compared against MusiteDeep, one of the most renowned tools for PTM prediction, using its phosphorylation (S,T) and phosphorylation (Y) modules. Acetyllysine is evaluated against DeepAcet, a robust model trained on over 20,000 acetylated protein sequences. For succinyllysine, the comparison is made with LMSuccSite, a model that incorporates recent trends and is based on language processing techniques. Methylarginine is compared against DeepRMethylSite, which combines CNN and LSTM architectures to enhance prediction capabilities. Methyllysine is measured against DeepKme, a model trained using a variety of protein data with different types of methylation. Due to the smaller data volumes and limited computational research on trimethyllysine and dimethyllysine, no comparative models are set for these modifications. The performance of UniTPM on independent testing data is assessed using the AUROC and AUPRC metrics, as illustrated in Figure 5. To further contextualize these results, the comparative statistics between UniTPM and the state-of-the-art model for each PTM type are presented in Table 2. Additional metrics, including accuracy, precision, recall, F1 score, and MCC, for evaluating UniPTM and its comparative models across various PTM types are detailed in Table 4. We speculate that UniPTM outperforms other models because it utilizes complete protein sequences during training, avoiding the loss of information critical for PTM formation. In contrast, traditional sliding window methods only consider the amino acids near the PTM sites, potentially overlooking broader sequence influences on PTM formation. In addition, before feeding protein sequences into the UniPTM model, we first apply pre-trained ESM-2 to embed them, ensuring that each amino acid is enriched with information from the entire protein sequence. The transformer encoder then further enhances these features, optimizing the model’s ability to capture and utilize complex biological data effectively. This approach allows us to score the PTM probability based solely on embeddings extracted from the target sites, which are not masked, unlike traditional models that depend on the information form adjacent amino acid residues of the target sites.

**Figure 5:**
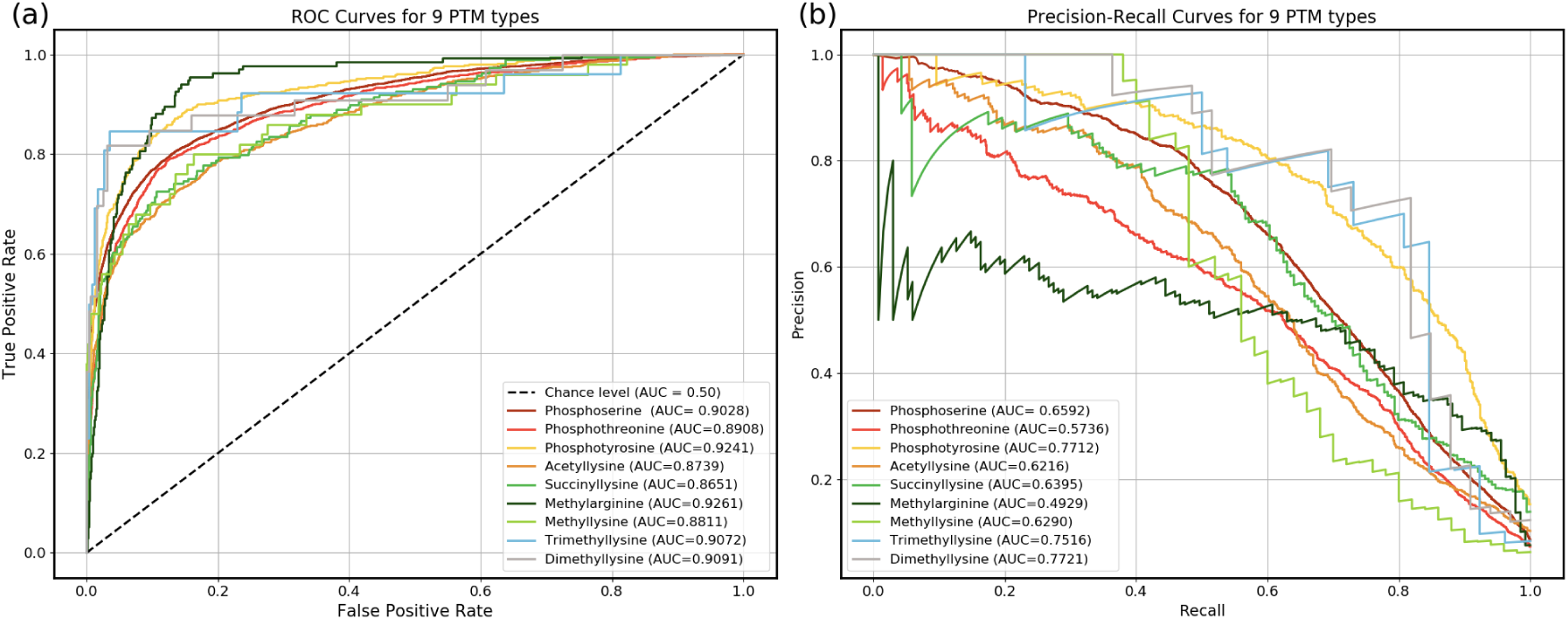
Performance of UniPTM on independent testing sets across 9 types of PTMs. (a) AUROCs. (b) AUPRCs.

**Table 2:** Comparison of PTM predictions for independent testing set by UniPTM and other models.

| PTM type | UniPTM |  | Existing model |  | Existing Model |
| --- | --- | --- | --- | --- | --- |
|  | AUROC | AUPRC | AUROC | AUPRC |  |
| Phosphoserine (S) | **0.9028** ± 0.0062 | **0.6592** ± 0.0176 | 0.8499 | 0.4278 | MusiteDeep (S,T)<sup>34</sup> |
| Phosphothreonine (T) | **0.8908** ± 0.0044 | **0.5736** ± 0.0310 | 0.8594 | 0.4378 | MusiteDeep (S,T)<sup>34</sup> |
| Phosphotyrosine (Y) | **0.9241** ± 0.0178 | **0.7712** ± 0.0397 | 0.9030 | 0.6487 | MusiteDeep (Y)<sup>34</sup> |
| Acetyllysine (K) | **0.8739** ± 0.0194 | **0.6216** ± 0.0720 | 0.6249 | 0.1214 | DeepAcet<sup>35</sup> |
| Succinyllysine (K) | **0.8651** ± 0.0274 | **0.6395** ± 0.0541 | 0.7102 | 0.2700 | LMSuccSite<sup>36</sup> |
| Methylarginine (R) | **0.9261** ± 0.0116 | **0.4929** ± 0.0540 | 0.9015 | 0.3180 | DeepRMethylSite<sup>37</sup> |
| Methyllysine (K) | **0.8811** ± 0.0579 | **0.6290** ± 0.1491 | 0.8100 | 0.2310 | DeepKme<sup>38</sup> |
| Trimethyllysine (K) | 0.9072 ± 0.0699 | 0.7516 ± 0.1394 | / | / | / |
| Dimethyllysine (K) | 0.9091 ± 0.0428 | 0.7721 ± 0.0761 | / | / | / |

As shown in Figure 5 and Table 2, UniPTM outperforms all seven comparative models on the independent test set. The AUC improvements range from a minimum of 2.28% for phosphotyrosine to a maximum of 28.48% for acetyllysine. The superior performance of UniPTM across multiple PTM types suggests that its approach to handling unsegmented, variable-length full-length protein sequences provides a significant advantage over traditional models that rely on processed fragments.

### Mapping of PTM predictions

The representations of PTM sites learned by UniPTM are visualized in this section to enhance the model’s interpretability. Each representation is mapped onto a 2D space using the dimension reduction tool t-SNE.^39^ T-SNE groups similar data points closer together and positions dissimilar data points farther apart. Unlike traditional methods, the data points in this work represent individual target amino acids embedded with the entire sequence information, rather than a protein fragment containing a single target amino acid residue. For example, the data points for phosphoserine consist of all phosphorylated serine residues plunked from full-length sequences, whereas non-phosphoserine data points comprise all the unphosphorylated serine residues from the protein sequences. Figure 6 represents features extracted by UniPTM architecture and the original site features for phosphoserine data. We can observe from Figure 6a that the majority of positive site samples are clustered a minor zone on the right side of the plot, with very few scattered among the negative site samples, indicating a clear separation. In Figure 6b, where the original site samples are embedded solely using ESM-2 model, show a slight clustering towards the lower right corner. Despite this, there is no clear distinction overall. The comparison of t-SNE plots for the other eight types of PTMs, as detailed in Figure S3-S5, aligns with the results observed for phospho-serine. The features of site samples generated by UniPTM model show significantly clearer clustering compared to those embedded with only pLM. By visualizing the representations generated by UniPTM, we demonstrate that the raw protein PTM sites can be transformed into biologically meaningful representations, thus enhancing the model’s interpretability.

**Figure 6:**
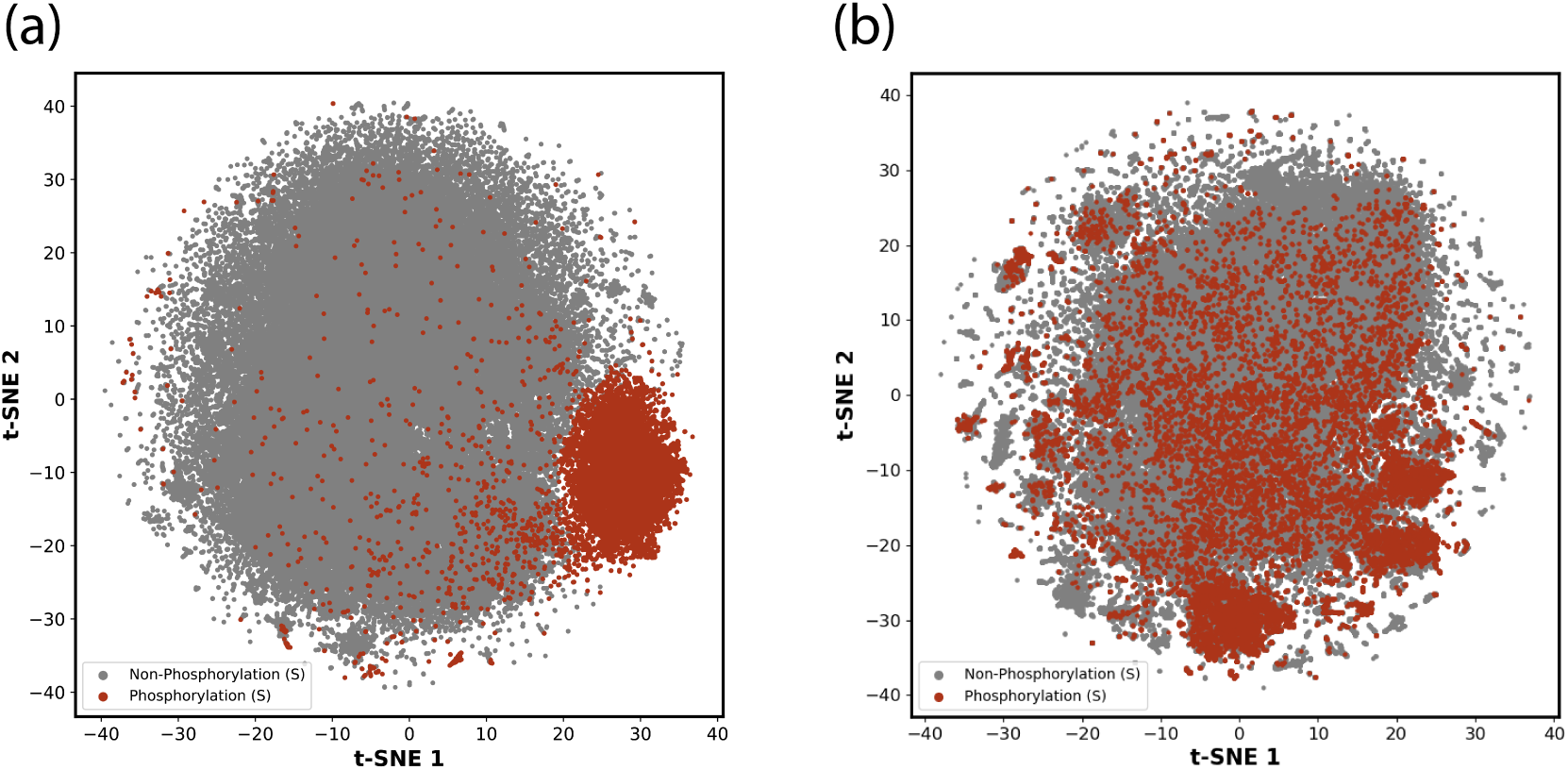
Visualization of features extracted by (a) UniPTM and (b) original site features by pre-trained ESM-2 model. Dark red dots represent positive site samples, which are the phosphorylated serine residues in full-length protein sequences, and gray dots represent negative site samples, which are non-phosphorylated serine residues.

## Discussion

In this study, we introduce UniPTM as a universal approach for PTM site predictions. UniPTM accepts and leverages full-length sequence data for both training and testing process. It has been shown to significantly outperform several other prominent tools across nine types of PTM on independent testing datasets. Especially, in the acetyllysine site prediction, UniPTM achieves over a 28.48% relative improvement in AUROC. After evaluating UniPTM, we attribute its promising performance primarily to three key factors: (1) UniPTM takes full-length protein sequence data as input, considering all amino acids across the entire protein that could influence the occurrence of PTMs, even if they are distal. (2) The pre-trained ESM-2 model embeds contextual information from the entire sequence into each residue, allowing every target amino acid to engage independently without loss of generality. (3) The customized loss function, TargetedBCELoss, masks all amino acids on a sequence except for the targeted amino acid, ensuring that only the sites with PTM potential participate in the loss calculation. This strategy achieves true site-level training, moving beyond the segment-level training typically seen in traditional slide-window based models. To our best knowledge, this is the first study where full-length protein sequences are adopted directly (without sliding window) for model training in the PTM annotation field.

In addition, we have established the PTMseq dataset, which consists of natural full-length protein sequences. This dataset includes the subsets of nine types of PTMed sequences, currently encompassing a total of 34,514 distinct PTM sites annotated across 12,203 full-length protein sequences. In future curation, we will perform updates every three months by literature and database synchronizations. Newly annotated sequences and new types of PTMs will be added to facilitate model training and benchmarking for other researchers in the community.

We do note, however, that future revisions to both our model and evaluation criteria will improve the utility of UniPTM. First and foremost, exploring newer or more complex deep learning architectures, such as Mamba,^40^ could potentially improve predictive performance. Second, packaging UniPTM as a user-friendly software tool or web server could facilitate accesses and uses by researchers who may not have extensive computational expertise. In addition, partnerships with experimental labs could provide real-world feedback and practical insights that refine the model’s accuracy and usability.

## Source code and dataset

The source code of UniPTM and the PTMseq dataset will be released after the manuscript has undergone peer review.

## References

(1) Consortium, U. UniProt: a worldwide hub of protein knowledge. Nucleic acids research 2019, 47, D506–D515.

(2) Meng, L.; Chan, W.-S.; Huang, L.; Liu, L.; Chen, X.; Zhang, W.; Wang, F.; Cheng, K.; Sun, H.; Wong, K.-C. Mini-review: Recent advances in post-translational modification site prediction based on deep learning. Computational and Structural Biotechnology Journal 2022,

(3) van Weeren, P. C.; de Bruyn, K. M.; de Vries-Smits, A. M.; Van Lint, J.; Boudewijn, M. Essential role for protein kinase B (PKB) in insulin-induced glycogen synthase kinase 3 inactivation: characterization of dominant-negative mutant of PKB. Journal of Biological Chemistry 1998, 273, 13150–13156.

(4) Verdin, E.; Ott, M. 50 years of protein acetylation: from gene regulation to epigenetics, metabolism and beyond. Nature reviews Molecular cell biology 2015, 16, 258–264.

(5) Dilworth, D.; Barsyte-Lovejoy, D. Targeting protein methylation: from chemical tools to precision medicines. Cellular and molecular life sciences 2019, 76, 2967–2985.

(6) Cohen, P. The origins of protein phosphorylation. Nature cell biology 2002, 4, E127– E130.

(7) Vlastaridis, P.; Kyriakidou, P.; Chaliotis, A.; Van de Peer, Y.; Oliver, S. G.; Amoutzias, G. D. Estimating the total number of phosphoproteins and phosphorylation sites in eukaryotic proteomes. Gigascience 2017, 6, giw015.

(8) Bannister, A.*; Miska, E. Regulation of gene expression by transcription factor acetylation. Cellular and Molecular Life Sciences CMLS 2000, 57, 1184–1192.

(9) Peng, C.; Wong, C. C. The story of protein arginine methylation: characterization, regulation, and function. Expert Review of Proteomics 2017, 14, 157–170.

(10) Wang, D.; Liu, D.; Yuchi, J.; He, F.; Jiang, Y.; Cai, S.; Li, J.; Xu, D. MusiteDeep: a deep-learning based webserver for protein post-translational modification site prediction and visualization. Nucleic Acids Research 2020, 48, W140–W146.

(11) Yu, Z.; Yu, J.; Wang, H.; Zhang, S.; Zhao, L.; Shi, S. PhosAF: An integrated deep learning architecture for predicting protein phosphorylation sites with AlphaFold2 predicted structures. Analytical Biochemistry 2024, 690, 115510.

(12) Li, Z.; Li, M.; Zhu, L.; Zhang, W. Improving PTM Site Prediction by Coupling of Multi-Granularity Structure and Multi-Scale Sequence Representation. arXiv preprint arXiv:2401.10211 2024,

(13) Peng, Z.; Schussheim, B.; Chatterjee, P. PTM-Mamba: A PTM-Aware Protein Language Model with Bidirectional Gated Mamba Blocks. bioRxiv 2024, 2024–02.

(14) Gutierrez, C. S.; Kassim, A. A.; Gutierrez, B. D.; Raines, R. T. Sitetack: A Deep Learning Model that Improves PTM Prediction by Using Known PTMs. bioRxiv 2024, 2024–06.

(15) Chou, K.-C. Prediction of signal peptides using scaled window. peptides 2001, 22, 1973–1979.

(16) Zhao, M.-X.; Chen, Q.; Li, F.; Fu, S.; Huang, B.; Zhao, Y. Protein phosphorylation database and prediction tools. Briefings in Bioinformatics 2023, 24, bbad090.

(17) Graves, D. J. Methods in Enzymology; Elsevier, 1983; Vol. 99; pp 268–278.

(18) Lin, Z.; Akin, H.; Rao, R.; Hie, B.; Zhu, Z.; Lu, W.; dos Santos Costa, A.; Fazel-Zarandi, M.; Sercu, T.; Candido, S.; others Language models of protein sequences at the scale of evolution enable accurate structure prediction. BioRxiv 2022, 2022, 500902.

(19) Huang, Y.; Niu, B.; Gao, Y.; Fu, L.; Li, W. CD-HIT Suite: a web server for clustering and comparing biological sequences. Bioinformatics 2010, 26, 680–682.

(20) Elnaggar, A.; Heinzinger, M.; Dallago, C.; Rihawi, G.; Wang, Y.; Jones, L.; Gibbs, T.; Feher, T.; Angerer, C.; Bhowmik, D.; Rost, B. ProtTrans: Towards Cracking the Lan-guage of Life’s Code Through Self-Supervised Deep Learning and High Performance Computing. bioRxiv 2020,

(21) Ahmed, E.; Heinzinger, M.; Dallago, C.; Rihawi, G.; Wang, Y.; Jones, L.; Gibbs, T.; Feher, T.; Angerer, C.; Martin, S.; others Prottrans: towards cracking the language of life’s code through self-supervised deep learning and high performance computing. bioRxiv 2020,

(22) Suzek, B. E.; Huang, H.; McGarvey, P.; Mazumder, R.; Wu, C. H. UniRef: comprehensive and non-redundant UniProt reference clusters. Bioinformatics 2007, 23, 1282–1288.

(23) Kenton, J. D. M.-W. C.; Toutanova, L. K. Bert: Pre-training of deep bidirectional transformers for language understanding. Proceedings of naacL-HLT. 2019; p 2.

(24) Meng, L.; Chen, X.; Cheng, K.; Chen, N.; Zheng, Z.; Wang, F.; Sun, H.; Wong, K.-C. TransPTM: a transformer-based model for non-histone acetylation site prediction. Briefings in Bioinformatics 2024, 25, bbae219.

(25) Maduranga, K. D. G.; Zadorozhnyy, V.; Ye, Q. Symmetry-structured convolutional neural networks. Neural Computing and Applications 2023, 35, 4421–4434.

(26) Torng, W.; Altman, R. B. 3D deep convolutional neural networks for amino acid environment similarity analysis. BMC bioinformatics 2017, 18, 1–23.

(27) Haviv, A.; Ram, O.; Press, O.; Izsak, P.; Levy, O. Transformer language models without positional encodings still learn positional information. arXiv preprint arXiv:2203.16634 2022,

(28) He, K.; Zhang, X.; Ren, S.; Sun, J. Deep residual learning for image recognition. Proceedings of the IEEE conference on computer vision and pattern recognition. 2016; pp 770–778.

(29) Ba, J. L.; Kiros, J. R.; Hinton, G. E. Layer normalization. arXiv 2016. arXiv preprint arXiv:1607.06450 2016, 1.

(30) Vaswani, A.; Shazeer, N.; Parmar, N.; Uszkoreit, J.; Jones, L.; Gomez, A. N.; Kaiser, Lt.; Polosukhin, I. Attention is all you need. Advances in neural information processing systems 2017, 30.

(31) Schwaller, P.; Laino, T.; Gaudin, T.; Bolgar, P.; Hunter, C. A.; Bekas, C.; Lee, A. A. Molecular transformer: a model for uncertainty-calibrated chemical reaction prediction. ACS central science 2019, 5, 1572–1583.

(32) Schwaller, P.; Probst, D.; Vaucher, A. C.; Nair, V. H.; Kreutter, D.; Laino, T.; Rey-mond, J.-L. Mapping the space of chemical reactions using attention-based neural networks. Nature machine intelligence 2021, 3, 144–152.

(33) Cao, Z.; Magar, R.; Wang, Y.; Barati Farimani, A. Moformer: self-supervised transformer model for metal–organic framework property prediction. Journal of the American Chemical Society 2023, 145, 2958–2967.

(34) Wang, D.; Zeng, S.; Xu, C.; Qiu, W.; Liang, Y.; Joshi, T.; Xu, D. MusiteDeep: a deep-learning framework for general and kinase-specific phosphorylation site prediction. Bioinformatics 2017, 33, 3909–3916.

(35) Wu, M.; Yang, Y.; Wang, H.; Xu, Y. A deep learning method to more accurately recall known lysine acetylation sites. BMC bioinformatics 2019, 20, 1–11.

(36) Pokharel, S.; Pratyush, P.; Heinzinger, M.; Newman, R. H.; Kc, D. B. Improving protein succinylation sites prediction using embeddings from protein language model. Scientific reports 2022, 12, 16933.

(37) Chaudhari, M.; Thapa, N.; Roy, K.; Newman, R. H.; Saigo, H.; Dukka, B. Deep-RMethylSite: a deep learning based approach for prediction of arginine methylation sites in proteins. Molecular omics 2020, 16, 448–454.

(38) Zou, G.; Zou, Y.; Ma, C.; Zhao, J.; Li, L. Development of an experiment-split method for benchmarking the generalization of a PTM site predictor: Lysine methylome as an example. PLoS Computational Biology 2021, 17, e1009682.

(39) Van der Maaten, L.; Hinton, G. Visualizing data using t-SNE. Journal of machine learning research 2008, 9.

(40) Gu, A.; Dao, T. Mamba: Linear-time sequence modeling with selective state spaces. arXiv preprint arXiv:2312.00752 2023,

