## Supporting information for "UniPTM: Multiple PTM site prediction on full-length protein sequence"

### Dataset construction details

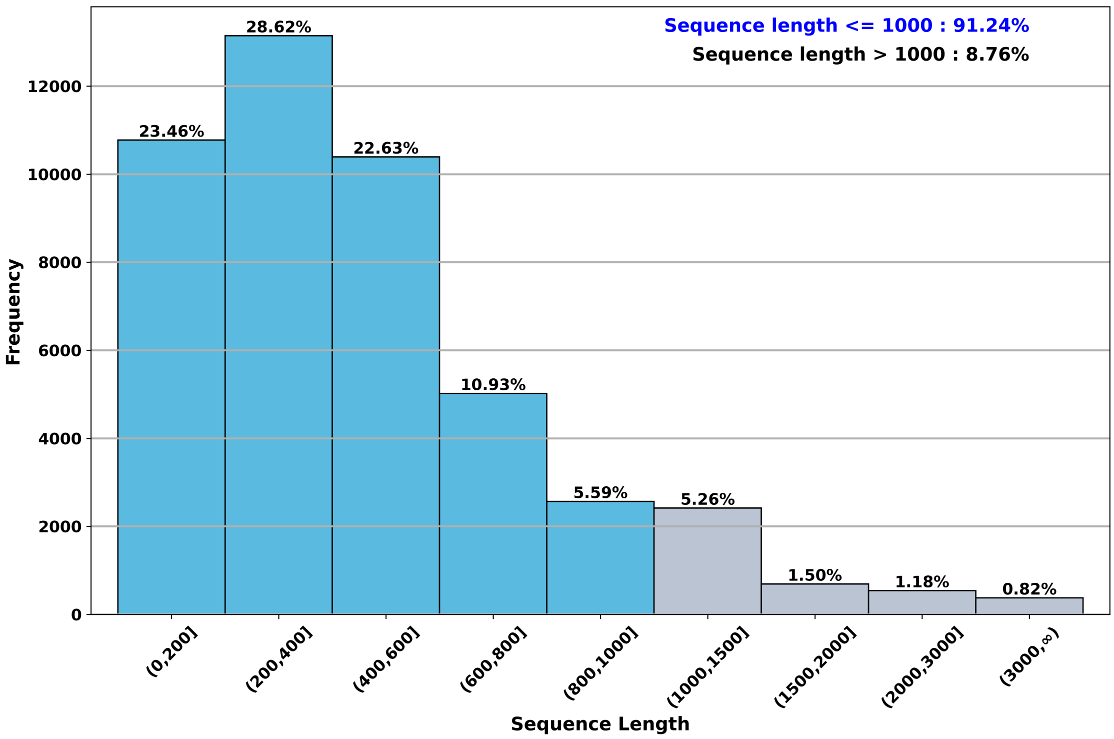


Figure S1: Distribution of sequence lengths in raw data.

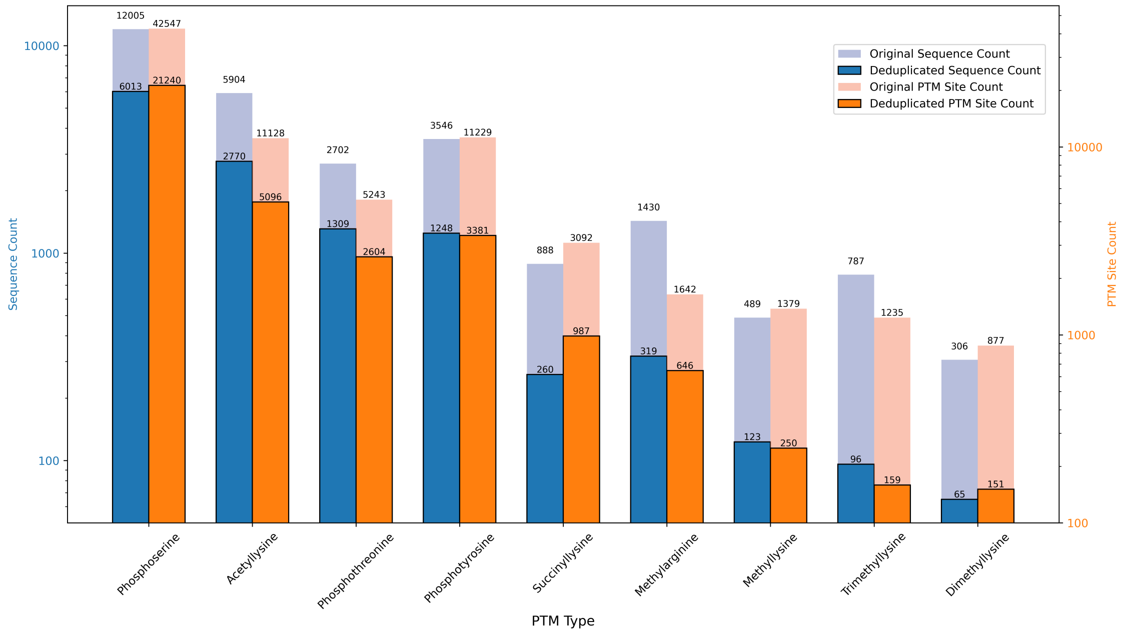


Figure S2: Data scale comparison in PTMseq pre- and post-CD-HIT deduplication processing.

Table S1: Results of 5-fold CV on entire data set before splitting.

| **One-hot** | | | | | | | |
|---|---|---|---|---|---|---|---|
| PTM type | Accuracy | Precision | Recall | F1 | MCC | AUROC | AUPRC |
| Phosphoserine | 0.9060 ± 0.0090 | 0.4659 ± 0.0354 | 0.4721 ± 0.0399 | 0.4671 ± 0.0155 | 0.4168 ± 0.0180 | 0.8483 ± 0.0057 | 0.4588 ± 0.0173 |
| Phosphothreonine | 0.9255 ± 0.0032 | 0.4556 ± 0.0395 | 0.4509 ± 0.0194 | 0.4526 ± 0.0260 | 0.4130 ± 0.0270 | 0.8531 ± 0.0070 | 0.4475 ± 0.0410 |
| Phosphotyrosine | 0.8994 ± 0.0297 | 0.6205 ± 0.1204 | 0.6090 ± 0.0319 | 0.6077 ± 0.0501 | 0.5548 ± 0.0660 | 0.8766 ± 0.0184 | 0.6481 ± 0.0720 |
| Acetyllysine | 0.9209 ± 0.0221 | 0.5766 ± 0.1414 | 0.4442 ± 0.0752 | 0.5001 ± 0.1021 | 0.4634 ± 0.1161 | 0.8197 ± 0.0368 | 0.5203 ± 0.1203 |
| Succinyllysine | 0.8238 ± 0.0358 | 0.3636 ± 0.1209 | 0.3228 ± 0.1290 | 0.3188 ± 0.0671 | 0.2348 ± 0.0783 | 0.7370 ± 0.0477 | 0.3438 ± 0.0378 |
| Methylarginine | 0.9235 ± 0.0148 | 0.3805 ± 0.0645 | 0.5027 ± 0.1203 | 0.4224 ± 0.0435 | 0.3932 ± 0.0409 | 0.8897 ± 0.0131 | 0.3573 ± 0.0742 |
| Methyllysine | 0.9477 ± 0.0127 | 0.6448 ± 0.2528 | 0.4288 ± 0.0748 | 0.5035 ± 0.1193 | 0.4945 ± 0.1405 | 0.8354 ± 0.0726 | 0.5108 ± 0.1801 |
| Trimethyllysine | 0.9547 ± 0.0193 | 0.6657 ± 0.3799 | 0.4495 ± 0.3246 | 0.5236 ± 0.3366 | 0.5257 ± 0.3310 | 0.8041 ± 0.1611 | 0.5677 ± 0.3061 |
| Dimethyllysine | 0.9462 ± 0.0199 | 0.7372 ± 0.0549 | 0.5923 ± 0.1405 | 0.6474 ± 0.0956 | 0.6281 ± 0.0867 | 0.8672 ± 0.0605 | 0.6901 ± 0.0805 |
| **ProtBert** | | | | | | | |
| PTM type | Accuracy | Precision | Recall | F1 | MCC | AUROC | AUPRC |
| Phosphoserine | 0.9342 ± 0.0076 | 0.6403 ± 0.0576 | 0.5890 ± 0.0570 | 0.6095 ± 0.0165 | 0.5768 ± 0.0168 | 0.8957 ± 0.0074 | 0.6426 ± 0.0183 |
| Phosphothreonine | 0.9301 ± 0.0161 | 0.5146 ± 0.1115 | 0.6017 ± 0.0530 | 0.5446 ± 0.0475 | 0.5152 ± 0.0459 | 0.8868 ± 0.0036 | 0.5679 ± 0.0376 |
| Phosphotyrosine | 0.9290 ± 0.0121 | 0.7183 ± 0.0475 | 0.7098 ± 0.0426 | 0.7137 ± 0.0418 | 0.6734 ± 0.0476 | 0.9227 ± 0.0122 | 0.7521 ± 0.0423 |
| Acetyllysine | 0.9356 ± 0.0090 | 0.6487 ± 0.0871 | 0.5813 ± 0.0351 | 0.6124 ± 0.0582 | 0.5789 ± 0.0641 | 0.8626 ± 0.0156 | 0.6267 ± 0.0552 |
| Succinyllysine | 0.9071 ± 0.0322 | 0.7171 ± 0.0993 | 0.5316 ± 0.0835 | 0.6084 ± 0.0802 | 0.5664 ± 0.0989 | 0.8547 ± 0.0437 | 0.6585 ± 0.0695 |
| Methylarginine | 0.9464 ± 0.0111 | 0.5359 ± 0.0732 | 0.5739 ± 0.0628 | 0.5484 ± 0.0295 | 0.5236 ± 0.0254 | 0.9235 ± 0.0086 | 0.4879 ± 0.0836 |
| Methyllysine | 0.9651 ± 0.0055 | 0.8228 ± 0.1419 | 0.5312 ± 0.1091 | 0.6418 ± 0.1093 | 0.6430 ± 0.1121 | 0.8812 ± 0.0628 | 0.6177 ± 0.1522 |
| Trimethyllysine | 0.9676 ± 0.0116 | 0.9031 ± 0.1216 | 0.6428 ± 0.1530 | 0.7360 ± 0.1115 | 0.7389 ± 0.0976 | 0.9021 ± 0.0585 | 0.7331 ± 0.1551 |
| Dimethyllysine | 0.9634 ± 0.0219 | 0.8984 ± 0.1334 | 0.6910 ± 0.0958 | 0.7744 ± 0.0840 | 0.7661 ± 0.0933 | 0.9282 ± 0.0297 | 0.8070 ± 0.0614 |

*Continued on next page*

Table S1 – *Continued from previous page*

| ProtT5 | | | | | | | |
|---|---|---|---|---|---|---|---|
| PTM type | Accuracy | Precision | Recall | F1 | MCC | AUROC | AUPRC |
| Phosphoserine | 0.9354 ± 0.0022 | 0.6367 ± 0.0296 | 0.6080 ± 0.0291 | 0.6212 ± 0.0146 | 0.5866 ± 0.0153 | 0.9032 ± 0.0059 | 0.6490 ± 0.0185 |
| Phosphothreonine | 0.9322 ± 0.0106 | 0.5121 ± 0.0945 | 0.5738 ± 0.0229 | 0.5383 ± 0.0542 | 0.5046 ± 0.0588 | 0.8888 ± 0.0100 | 0.5358 ± 0.0612 |
| Phosphotyrosine | 0.9206 ± 0.0111 | 0.6606 ± 0.0530 | 0.7513 ± 0.0269 | 0.7022 ± 0.0360 | 0.6590 ± 0.0391 | 0.9180 ± 0.0117 | 0.7522 ± 0.0366 |
| Acetyllysine | 0.9387 ± 0.0058 | 0.6738 ± 0.0481 | 0.5871 ± 0.0538 | 0.6256 ± 0.0370 | 0.5951 ± 0.0365 | 0.8816 ± 0.0183 | 0.6392 ± 0.0490 |
| Succinyllysine | 0.8971 ± 0.0243 | 0.6435 ± 0.0467 | 0.5358 ± 0.0886 | 0.5812 ± 0.0606 | 0.5284 ± 0.0691 | 0.8589 ± 0.0356 | 0.6363 ± 0.0484 |
| Methylarginine | 0.9440 ± 0.0126 | 0.5138 ± 0.0531 | 0.5793 ± 0.0784 | 0.5399 ± 0.0384 | 0.5139 ± 0.0363 | 0.9308 ± 0.0061 | 0.5135 ± 0.0613 |
| Methyllysine | 0.9646 ± 0.0060 | 0.7843 ± 0.1290 | 0.5544 ± 0.1264 | 0.6465 ± 0.1188 | 0.6407 ± 0.1207 | 0.8850 ± 0.0599 | 0.6272 ± 0.1686 |
| Trimethyllysine | 0.9705 ± 0.0139 | 0.9091 ± 0.1107 | 0.6759 ± 0.1696 | 0.7623 ± 0.1230 | 0.7632 ± 0.1143 | 0.9039 ± 0.0719 | 0.7491 ± 0.1515 |
| Dimethyllysine | 0.9611 ± 0.0239 | 0.8458 ± 0.0887 | 0.7008 ± 0.1276 | 0.7613 ± 0.0946 | 0.7472 ± 0.1019 | 0.9082 ± 0.0356 | 0.7853 ± 0.0821 |
| **ESM-2** | | | | | | | |
| PTM type | Accuracy | Precision | Recall | F1 | MCC | AUROC | AUPRC |
| Phosphoserine | 0.9372 ± 0.0031 | 0.6513 ± 0.0301 | 0.6046 ± 0.0248 | 0.6265 ± 0.0170 | 0.5931 ± 0.0183 | 0.9019 ± 0.0061 | 0.6634 ± 0.0203 |
| Phosphothreonine | 0.9264 ± 0.0070 | 0.4727 ± 0.0174 | 0.6338 ± 0.0279 | 0.5411 ± 0.0127 | 0.5086 ± 0.0155 | 0.8944 ± 0.0101 | 0.5592 ± 0.0191 |
| Phosphotyrosine | 0.9321 ± 0.0062 | 0.7287 ± 0.0524 | 0.7249 ± 0.0396 | 0.7258 ± 0.0352 | 0.6877 ± 0.0376 | 0.9229 ± 0.0160 | 0.7694 ± 0.0457 |
| Acetyllysine | 0.9400 ± 0.0054 | 0.6814 ± 0.0308 | 0.5837 ± 0.0703 | 0.6277 ± 0.0532 | 0.5980 ± 0.0519 | 0.8709 ± 0.0200 | 0.6295 ± 0.0628 |
| Succinyllysine | 0.9087 ± 0.0223 | 0.7087 ± 0.0484 | 0.5567 ± 0.0942 | 0.6188 ± 0.0635 | 0.5766 ± 0.0665 | 0.8783 ± 0.0327 | 0.6764 ± 0.0499 |
| Methylarginine | 0.9417 ± 0.0145 | 0.5156 ± 0.1095 | 0.5824 ± 0.1109 | 0.5310 ± 0.0297 | 0.5107 ± 0.0267 | 0.9239 ± 0.0059 | 0.4831 ± 0.0686 |
| Methyllysine | 0.9639 ± 0.0099 | 0.7696 ± 0.1810 | 0.5577 ± 0.1305 | 0.6443 ± 0.1435 | 0.6362 ± 0.1510 | 0.8769 ± 0.0660 | 0.6186 ± 0.1817 |
| Trimethyllysine | 0.9670 ± 0.0118 | 0.8711 ± 0.1143 | 0.6619 ± 0.1645 | 0.7360 ± 0.1129 | 0.7348 ± 0.0982 | 0.9073 ± 0.0715 | 0.7437 ± 0.1358 |
| Dimethyllysine | 0.9606 ± 0.0246 | 0.8793 ± 0.1351 | 0.6862 ± 0.1296 | 0.7611 ± 0.0970 | 0.7521 ± 0.1056 | 0.9015 ± 0.0715 | 0.7781 ± 0.0882 |

#### Transformer model and multi-head attention

In this research, we apply the encoder component of the transformer model as the foundation for the UniPTM framework. Transformers are a category of deep learning models that have made significant advancements in natural language processing (NLP)[1,2] and have recently been utilized to model protein sequences.[3] The encoder component learns the latent representation of the input sequence through the self-attention mechanism. The self-attention mechanism has been used in conjunction with sequential models, such as recurrent neural networks (RNN)[4] and LSTM,[5] to address declines in model performance when processing long sequences. The transformer model discards the recurrent architecture found in earlier sequential models and depends entirely on the self-attention mechanism to learn input sequence‘s representations. The benefits of this model architecture are various. Self-attention reduces the computational complexity per layer, allowing for faster training as the model relies solely on self-attention, which primarily involves matrix multiplications. Additionally, the self-attention mechanism effectively facilitates learning long-range dependencies within the input sequence.

The weights used for linearly projecting the entire input sequence are represented by the matrices $W_q$, $W_k$, and $W_v$. In this research, we applied the scaled dot-product attention:

$$\text{Attention}(Q, K, V) = \text{softmax}\left(\frac{QK^\top}{\sqrt{d_k}}\right)V \tag{1}$$

Multi-head attention involves using several linear projection matrices simultaneously to compute attention. In multi-head attention, the matrices $W_q$, $W_k$, and $W_v$ are initialized differently for each head, allowing the model to capture various representations of the input sequence from different subspaces. If we assume that there are $h$ attention heads, the multi-head attention can be computed using the following formula:

$$\text{Multi-Head}(Q, K, V) = \text{Concat}(\text{head}_1, \text{head}_2, \ldots, \text{head}_h)W^O \tag{2}$$

where each $head_i$ is computed as:

$$\text{head}_i = \text{Attention}(QW_i^Q, KW_i^K, VW_i^V) \tag{3}$$

Here, $W_i^Q$, $W_i^K$, and $W_i^V$ are the parameter matrices for the $i$-th head; and $W^O$ is the output projection matrix that combines the outputs of all heads. In this study, $d_{\text{emb}} = 256$ and the number of head $h = 8$. The dimension of the $W_i^Q$, $W_i^K$, and $W_v^i$ linear projection matrices is $W_i^Q \in \mathbb{R}^{d_{emb} \times d_k}$, $W_i^Q \in \mathbb{R}^{d_{emb} \times d_k}$, and $W_i^V \in \mathbb{R}^{d_{emb} \times d_k}$, where $d_{\text{emb}} = 256$ and $d_v = d_k = d_{emb}/h = 32$. Moreover, $W^O \in \mathbb{R}^{d_{emb} \times d_{emb}}$.

### Training details

In this section, we provide detailed descriptions of the hyperparameters (Table S1) used for training the UniPTM model, along with the equations (C.4-C.8) for the five evaluation criteria.

Table S2: Hyperparameters for the UniPTM model training

| Batch Size | lr | Optimizer | Epochs | Emb size | Train/Val | Weight Decay | dropout rate | pos_weight |
|---|---|---|---|---|---|---|---|---|
| 32 | 5e-5 | Adam | 200 | 1024/1280 | 0.9/0.1 | 1e-5 | 0.5 | 3 |

$$\text{Accuracy} = \frac{TP + TN}{TP + TN + FP + FN} \tag{4}$$

$$\text{Precision} = \frac{TP}{TP + FP} \tag{5}$$

$$\text{Recall} = \frac{TP}{TP + FN} \tag{6}$$

$$\text{F1 score} = \frac{2 \times \text{ Precision } \times \text{ Recall}}{\text{Precision } + \text{ Recall}} \tag{7}$$

$$\text{MCC} = \frac{TP \cdot TN - FP \cdot FN}{\sqrt{(TP+FP)(TP+FN)(TN+FP)(TN+FN)}} \tag{8}$$

### Model evaluation results

Table S3: Results of 5-fold CV on training data.

| **One-hot** | | | | | | | |
|---|---|---|---|---|---|---|---|
| PTM type | Accuracy | Precision | Recall | F1 | MCC | AUROC | AUPRC |
| Phosphoserine | 0.9076 ± 0.0081 | 0.4712 ± 0.0396 | 0.4168 ± 0.0233 | 0.4419 ± 0.0279 | 0.3928 ± 0.0328 | 0.8367 ± 0.0078 | 0.4367 ± 0.0353 |
| Phosphothreonine | 0.9176 ± 0.0123 | 0.3953 ± 0.0686 | 0.3889 ± 0.0583 | 0.3916 ± 0.0618 | 0.3477 ± 0.0675 | 0.8298 ± 0.0172 | 0.3751 ± 0.0764 |
| Phosphotyrosine | 0.8838 ± 0.0190 | 0.5341 ± 0.0744 | 0.5735 ± 0.0521 | 0.5513 ± 0.0536 | 0.4863 ± 0.0629 | 0.8482 ± 0.0200 | 0.5749 ± 0.0695 |
| Acetyllysine | 0.9039 ± 0.0103 | 0.4367 ± 0.0488 | 0.3863 ± 0.0581 | 0.4085 ± 0.0470 | 0.3581 ± 0.0521 | 0.7912 ± 0.0229 | 0.4088 ± 0.0601 |
| Succinyllysine | 0.8431 ± 0.0381 | 0.1963 ± 0.1795 | 0.1789 ± 0.1755 | 0.1846 ± 0.1735 | 0.1271 ± 0.1182 | 0.6601 ± 0.0838 | 0.2644 ± 0.0711 |
| Methylarginine | 0.9276 ± 0.0167 | 0.3918 ± 0.0750 | 0.4383 ± 0.0653 | 0.4074 ± 0.0467 | 0.3736 ± 0.0472 | 0.8702 ± 0.0200 | 0.3519 ± 0.0909 |
| Methyllysine | 0.9443 ± 0.0161 | 0.5517 ± 0.3704 | 0.3241 ± 0.1880 | 0.4007 ± 0.2431 | 0.3982 ± 0.2509 | 0.8119 ± 0.0529 | 0.4908 ± 0.1251 |
| Trimethyllysine | 0.9516 ± 0.0236 | 0.6795 ± 0.3815 | 0.4260 ± 0.2419 | 0.5219 ± 0.2926 | 0.5214 ± 0.2921 | 0.8309 ± 0.1145 | 0.6348 ± 0.1680 |
| Dimethyllysine | 0.9348 ± 0.0254 | 0.7324 ± 0.4237 | 0.2807 ± 0.2282 | 0.3776 ± 0.2714 | 0.4154 ± 0.2614 | 0.7980 ± 0.1060 | 0.5633 ± 0.1335 |
| **ProtBert** | | | | | | | |
| PTM type | Accuracy | Precision | Recall | F1 | MCC | AUROC | AUPRC |
| Phosphoserine | 0.9331 ± 0.0039 | 0.6416 ± 0.0306 | 0.5420 ± 0.0262 | 0.5867 ± 0.0090 | 0.5534 ± 0.0100 | 0.8837 ± 0.0043 | 0.6126 ± 0.0141 |
| Phosphothreonine | 0.9331 ± 0.0069 | 0.5088 ± 0.0471 | 0.5132 ± 0.0426 | 0.5101 ± 0.0370 | 0.4747 ± 0.0402 | 0.8744 ± 0.0045 | 0.5048 ± 0.0494 |
| Phosphotyrosine | 0.9108 ± 0.0052 | 0.6250 ± 0.0436 | 0.6994 ± 0.0219 | 0.6595 ± 0.0292 | 0.6101 ± 0.0307 | 0.9097 ± 0.0160 | 0.7076 ± 0.0286 |
| Acetyllysine | 0.9278 ± 0.0081 | 0.5985 ± 0.0547 | 0.5059 ± 0.0605 | 0.5461 ± 0.0472 | 0.5108 ± 0.0493 | 0.8348 ± 0.0234 | 0.5595 ± 0.0491 |
| Succinyllysine | 0.8961 ± 0.0286 | 0.7613 ± 0.1427 | 0.3465 ± 0.1177 | 0.4609 ± 0.1217 | 0.4582 ± 0.0978 | 0.8154 ± 0.0371 | 0.5647 ± 0.0645 |
| Methylarginine | 0.9456 ± 0.0125 | 0.5239 ± 0.0623 | 0.4813 ± 0.1122 | 0.4969 ± 0.0784 | 0.4714 ± 0.0788 | 0.9028 ± 0.0235 | 0.4331 ± 0.0941 |
| Methyllysine | 0.9575 ± 0.0108 | 0.7642 ± 0.1417 | 0.4957 ± 0.0836 | 0.6006 ± 0.1020 | 0.5948 ± 0.1116 | 0.8683 ± 0.0507 | 0.5923 ± 0.1395 |
| Trimethyllysine | 0.9678 ± 0.0163 | 0.8800 ± 0.1460 | 0.6613 ± 0.1582 | 0.7443 ± 0.1138 | 0.7418 ± 0.1182 | 0.8865 ± 0.0725 | 0.7551 ± 0.1448 |
| Dimethyllysine | 0.9553 ± 0.0200 | 0.8556 ± 0.1089 | 0.6013 ± 0.1882 | 0.6819 ± 0.1241 | 0.6831 ± 0.0954 | 0.8566 ± 0.0927 | 0.7122 ± 0.1416 |

*Continued on next page*

Table S3 – *Continued from previous page*

| **ProtT5** | | | | | | | |
|---|---|---|---|---|---|---|---|
| PTM type | Accuracy | Precision | Recall | F1 | MCC | AUROC | AUPRC |
| Phosphoserine | 0.9329 ± 0.0063 | 0.6403 ± 0.0577 | 0.5619 ± 0.0491 | 0.5948 ± 0.0035 | 0.5621 ± 0.0068 | 0.8923 ± 0.0043 | 0.6231 ± 0.0068 |
| Phosphothreonine | 0.9327 ± 0.0101 | 0.5118 ± 0.0756 | 0.5944 ± 0.0306 | 0.5466 ± 0.0383 | 0.5143 ± 0.0406 | 0.8893 ± 0.0033 | 0.5489 ± 0.0487 |
| Phosphotyrosine | 0.9127 ± 0.0078 | 0.6298 ± 0.0318 | 0.7089 ± 0.0584 | 0.6665 ± 0.0401 | 0.6182 ± 0.0444 | 0.9073 ± 0.0127 | 0.7001 ± 0.0479 |
| Acetyllysine | 0.9210 ± 0.0045 | 0.5476 ± 0.0451 | 0.5247 ± 0.0546 | 0.5325 ± 0.0211 | 0.4917 ± 0.0202 | 0.8483 ± 0.0188 | 0.5364 ± 0.0318 |
| Succinyllysine | 0.8838 ± 0.0265 | 0.6379 ± 0.1527 | 0.4082 ± 0.1498 | 0.4674 ± 0.1127 | 0.4335 ± 0.0704 | 0.8218 ± 0.0415 | 0.5560 ± 0.0692 |
| Methylarginine | 0.9483 ± 0.0084 | 0.5475 ± 0.0666 | 0.4581 ± 0.0781 | 0.4976 ± 0.0712 | 0.4734 ± 0.0724 | 0.9109 ± 0.0275 | 0.4877 ± 0.0895 |
| Methyllysine | 0.9559 ± 0.0097 | 0.7644 ± 0.1451 | 0.4689 ± 0.0647 | 0.5791 ± 0.0844 | 0.5768 ± 0.0957 | 0.8551 ± 0.0446 | 0.5865 ± 0.1310 |
| Trimethyllysine | 0.9676 ± 0.0139 | 0.8942 ± 0.0769 | 0.6381 ± 0.1504 | 0.7341 ± 0.0915 | 0.7349 ± 0.0847 | 0.8994 ± 0.0791 | 0.7504 ± 0.1378 |
| Dimethyllysine | 0.9418 ± 0.0232 | 0.8548 ± 0.1068 | 0.4266 ± 0.2142 | 0.5293 ± 0.1838 | 0.5551 ± 0.1295 | 0.8327 ± 0.1040 | 0.6357 ± 0.1412 |
| **ESM-2** | | | | | | | |
| PTM type | Accuracy | Precision | Recall | F1 | MCC | AUROC | AUPRC |
| Phosphoserine | 0.9338 ± 0.0055 | 0.6472 ± 0.0578 | 0.5597 ± 0.0486 | 0.5968 ± 0.0109 | 0.5647 ± 0.0117 | 0.8894 ± 0.0050 | 0.6291 ± 0.0094 |
| Phosphothreonine | 0.9310 ± 0.0090 | 0.4981 ± 0.0524 | 0.5884 ± 0.0360 | 0.5375 ± 0.0293 | 0.5038 ± 0.0320 | 0.8846 ± 0.0060 | 0.5373 ± 0.0419 |
| Phosphotyrosine | 0.9198 ± 0.0074 | 0.6736 ± 0.0436 | 0.6943 ± 0.0615 | 0.6812 ± 0.0209 | 0.6372 ± 0.0225 | 0.9150 ± 0.0181 | 0.7340 ± 0.0367 |
| Acetyllysine | 0.9320 ± 0.0070 | 0.6452 ± 0.0248 | 0.4793 ± 0.0711 | 0.5467 ± 0.0426 | 0.5193 ± 0.0356 | 0.8491 ± 0.0184 | 0.5580 ± 0.0365 |
| Succinyllysine | 0.8989 ± 0.0264 | 0.6973 ± 0.1027 | 0.4304 ± 0.0749 | 0.5289 ± 0.0688 | 0.4950 ± 0.0803 | 0.8449 ± 0.0323 | 0.5950 ± 0.0755 |
| Methylarginine | 0.9459 ± 0.0074 | 0.5199 ± 0.0640 | 0.5089 ± 0.0929 | 0.5110 ± 0.0664 | 0.4844 ± 0.0683 | 0.9016 ± 0.0286 | 0.4566 ± 0.1121 |
| Methyllysine | 0.9578 ± 0.0091 | 0.7661 ± 0.1249 | 0.4991 ± 0.0733 | 0.6035 ± 0.0880 | 0.5976 ± 0.0961 | 0.8521 ± 0.0572 | 0.5776 ± 0.1582 |
| Trimethyllysine | 0.9687 ± 0.0174 | 0.9206 ± 0.0865 | 0.6571 ± 0.1528 | 0.7535 ± 0.0733 | 0.7565 ± 0.0689 | 0.8999 ± 0.0724 | 0.7600 ± 0.1390 |
| Dimethyllysine | 0.9536 ± 0.0176 | 0.8350 ± 0.1073 | 0.5936 ± 0.1680 | 0.6745 ± 0.0974 | 0.6724 ± 0.0816 | 0.8468 ± 0.0814 | 0.6701 ± 0.1064 |

Table S4: Performance comparison of state-of-the-art models and UniPTM on independent testing set.

| PTM type | Accuracy | Precision | Recall | F1 | MCC | AUROC | AUPRC |
|---|---|---|---|---|---|---|---|
| Phosphoserine | 0.9389 ± 0.0015 | 0.6691 ± 0.0157 | 0.5910 ± 0.0197 | 0.6275 ± 0.0141 | 0.5958 ± 0.0143 | 0.9028 ± 0.0062 | 0.6592 ± 0.0176 |
| MusiteDeep (S,T) | 0.7528 | 0.2272 | 0.7915 | 0.3531 | 0.3305 | 0.8399 | 0.4278 |
| Phosphothreonine | 0.9385 ± 0.0048 | 0.5522 ± 0.0536 | 0.5827 ± 0.0461 | 0.5640 ± 0.0240 | 0.5330 ± 0.0234 | 0.8908 ± 0.0044 | 0.5736 ± 0.0310 |
| MusiteDeep (S,T) | 0.8901 | 0.3394 | 0.5884 | 0.4305 | 0.3919 | 0.8594 | 0.4378 |
| Phosphotyrosine | 0.9299 ± 0.0097 | 0.7127 ± 0.0596 | 0.7361 ± 0.0369 | 0.7230 ± 0.0383 | 0.6838 ± 0.0418 | 0.9241 ± 0.0178 | 0.7712 ± 0.0397 |
| MusiteDeep (Y) | 0.7843 | 0.3591 | 0.8772 | 0.5096 | 0.4668 | 0.9030 | 0.6487 |
| Acetyllysine | 0.9358 ± 0.0109 | 0.6500 ± 0.1010 | 0.5902 ± 0.0402 | 0.6170 ± 0.0658 | 0.5839 ± 0.0726 | 0.8739 ± 0.0194 | 0.6216 ± 0.0720 |
| DeepAcet | 0.5040 | 0.1081 | 0.7111 | 0.1877 | 0.1074 | 0.6249 | 0.1214 |
| Succinyllysine | 0.9019 ± 0.0242 | 0.6774 ± 0.0590 | 0.5328 ± 0.1071 | 0.5902 ± 0.0719 | 0.5447 ± 0.0756 | 0.8651 ± 0.0274 | 0.6395 ± 0.0541 |
| LMSuccSite | 0.4365 | 0.1854 | 0.8918 | 0.3070 | 0.1880 | 0.7102 | 0.2700 |
| Methylarginine | 0.9336 ± 0.0196 | 0.4645 ± 0.0880 | 0.6083 ± 0.1307 | 0.5111 ± 0.0250 | 0.4909 ± 0.0213 | 0.9261 ± 0.0116 | 0.4929 ± 0.0540 |
| DeepRMethylSite | 0.9191 | 0.2917 | 0.5283 | 0.3758 | 0.3534 | 0.9015 | 0.3108 |
| Methyllysine | 0.9603 ± 0.0117 | 0.7326 ± 0.2043 | 0.5467 ± 0.1134 | 0.6217 ± 0.1409 | 0.6108 ± 0.1514 | 0.8811 ± 0.0579 | 0.6290 ± 0.1491 |
| DeepKme | 0.6030 | 0.1380 | 0.7910 | 0.2160 | 0.2240 | 0.8100 | 0.2310 |
| Trimethyllysine | 0.9681 ± 0.0096 | 0.8771 ± 0.0931 | 0.6729 ± 0.1321 | 0.7501 ± 0.0758 | 0.7468 ± 0.0657 | 0.9072 ± 0.0699 | 0.7516 ± 0.1394 |
| Dimethyllysine | 0.9586 ± 0.0247 | 0.8465 ± 0.1223 | 0.6896 ± 0.1144 | 0.7518 ± 0.0842 | 0.7387 ± 0.0941 | 0.9091 ± 0.0428 | 0.7721 ± 0.0761 |

#### UniPTM mechanism visualization

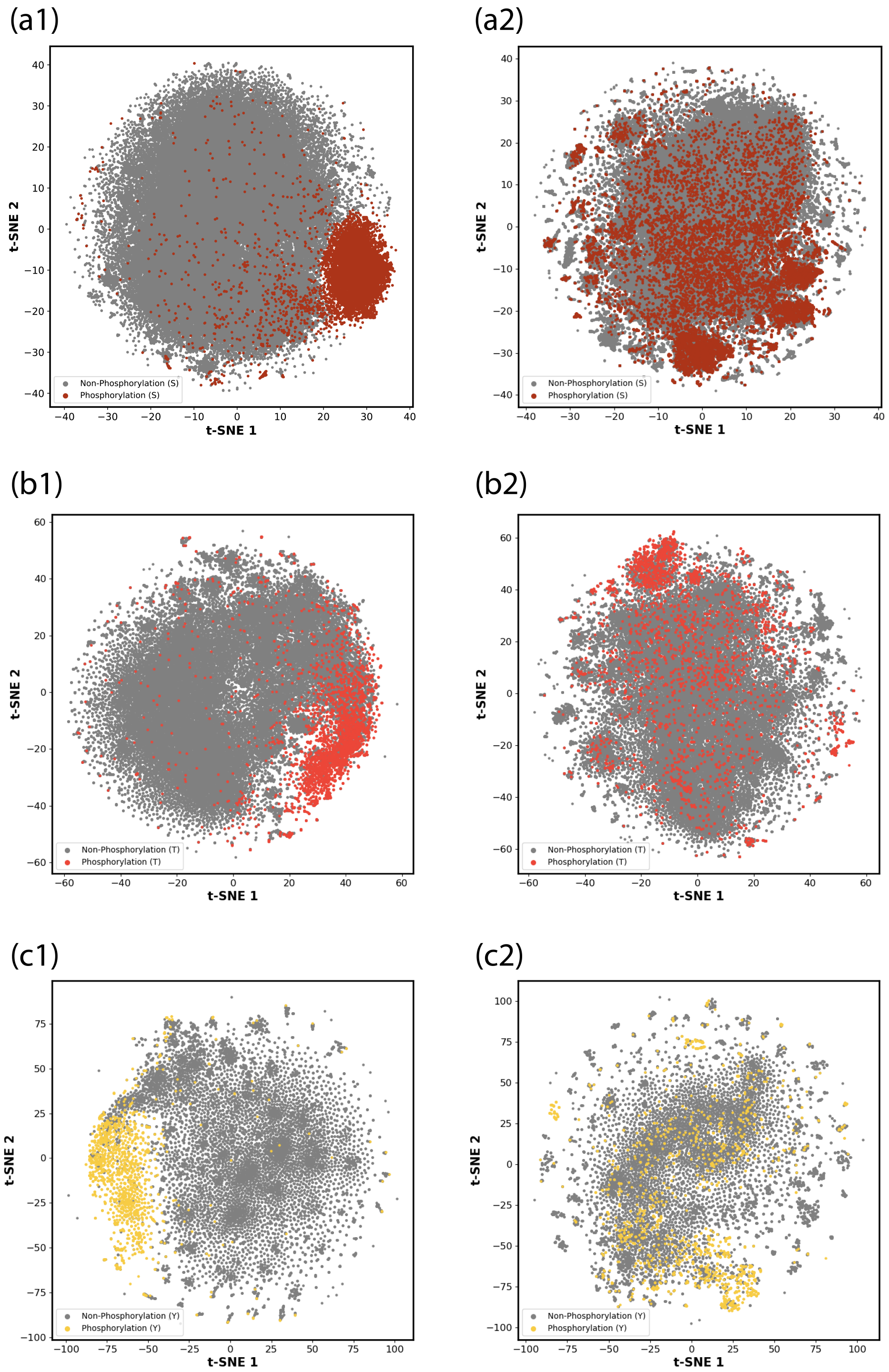


Figure S3: Visualization of abstract features extracted by UniPTM and original site features by pre-trained ESM-2 model (Part I: phosphoserine, phosphothreosine, and phosphotyrosine). Colored dots represent positive site samples, which are the PTM residues in full-length protein sequences, and gray dots represent negative site samples, which are non-PTM residues.

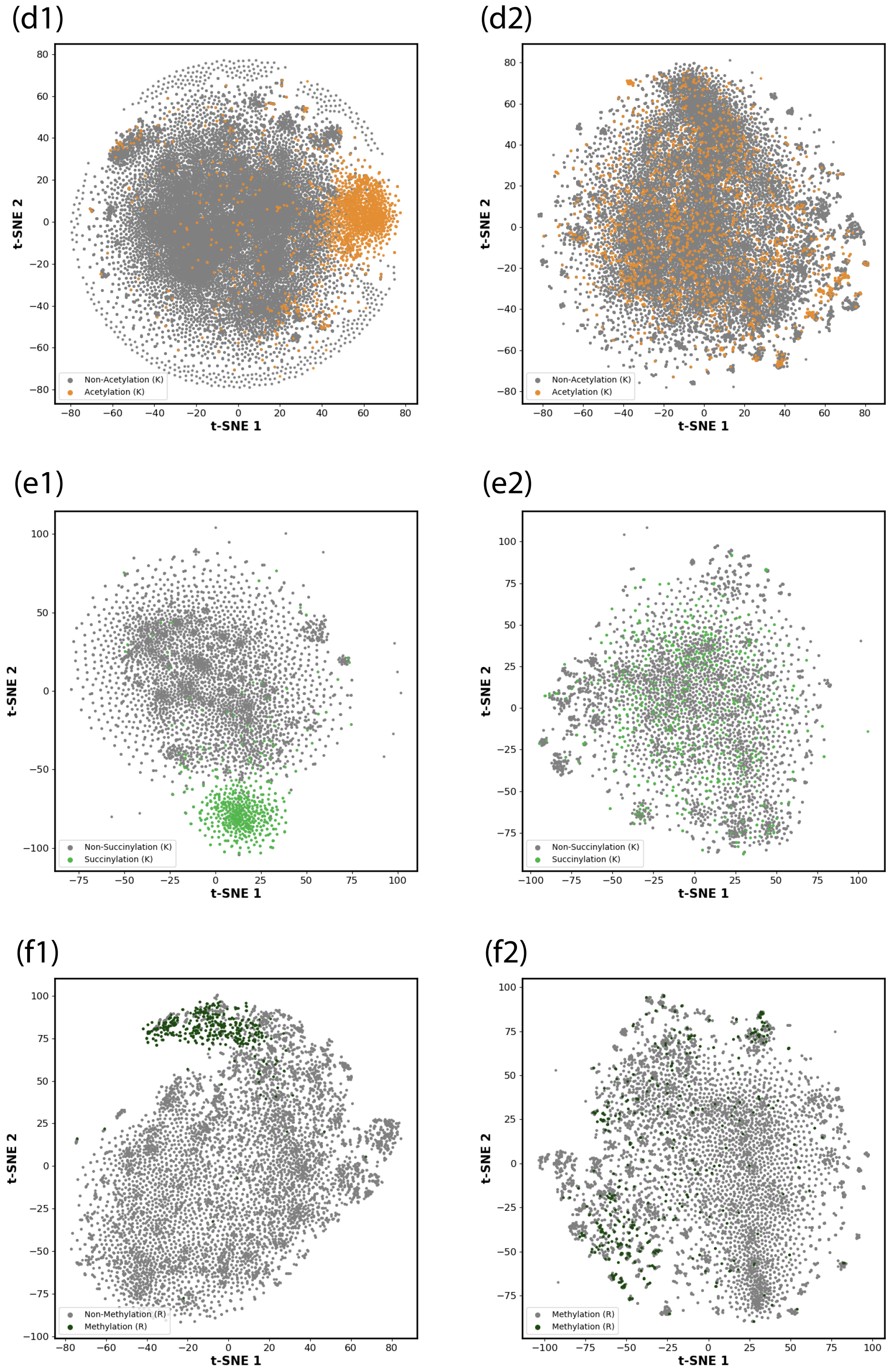


Figure S4: Visualization of abstract features extracted by UniPTM and original site features by pre-trained ESM-2 model (Part II: acetyllysine, succinyllysine, and methylarginine). Colored dots represent positive site samples, which are the PTM residues in full-length protein sequences, and gray dots represent negative site samples, which are non-PTM residues.

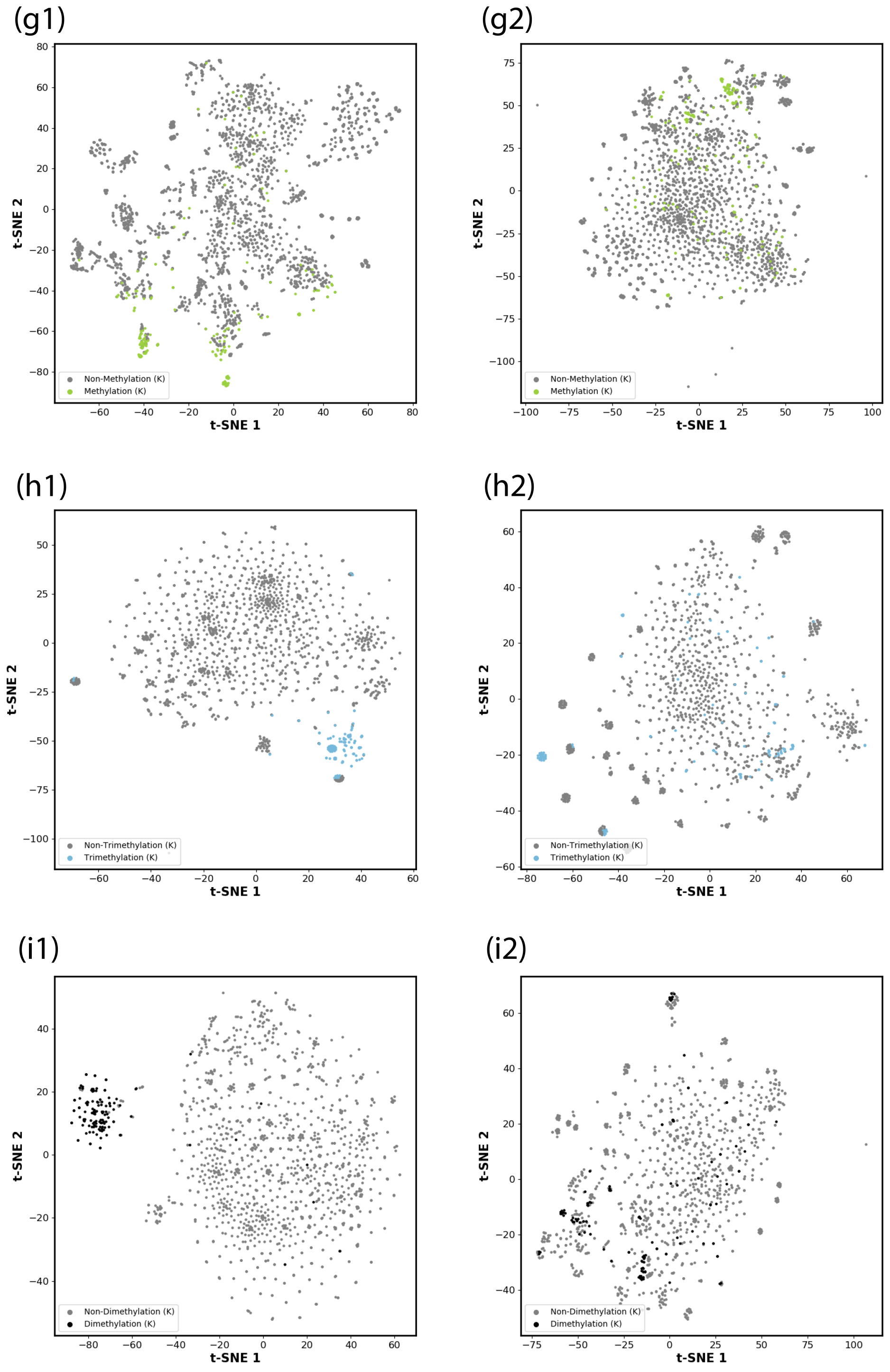


Figure S5: Visualization of abstract features extracted by UniPTM and original site features by pre-trained ESM-2 model (Part III: methyllysine, trimethyllysine, and dimethyllysine). Colored dots represent positive site samples, which are the PTM residues in full-length protein sequences, and gray dots represent negative site samples, which are non-PTM residues.

### References

(1) Vaswani, A.; Shazeer, N.; Parmar, N.; Uszkoreit, J.; Jones, L.; Gomez, A. N.; Kaiser, Ł.; Polosukhin, I. Attention is all you need. *Advances in neural information processing systems* **2017**, *30*.

(2) Brown, T.; Mann, B.; Ryder, N.; Subbiah, M.; Kaplan, J. D.; Dhariwal, P.; Neelakantan, A.; Shyam, P.; Sastry, G.; Askell, A.; others Language models are few-shot learners. *Advances in neural information processing systems* **2020**, *33*, 1877–1901.

(3) Chowdhury, R.; Bouatta, N.; Biswas, S.; Floristean, C.; Kharkar, A.; Roy, K.; Rochereau, C.; Ahdritz, G.; Zhang, J.; Church, G. M.; others Single-sequence protein structure prediction using a language model and deep learning. *Nature Biotechnology* **2022**, *40*, 1617–1623.

(4) Bahdanau, D.; Cho, K.; Bengio, Y. Neural machine translation by jointly learning to align and translate. *arXiv preprint arXiv:1409.0473* **2014**,

(5) Hochreiter, S.; Schmidhuber, J. Long short-term memory. *Neural computation* **1997**, *9*, 1735–1780.
